# Mining the human gut microbiome identifies mycobacterial d-arabinan degrading enzymes

**DOI:** 10.1101/2022.07.22.500997

**Authors:** Omar Al-Jourani, Samuel Benedict, Jennifer Ross, Abigail Layton, Phillip van der Peet, Victoria M. Marando, Nicholas P. Bailey, Tiaan Heunis, Joseph Manion, Francesca Mensitieri, Aaron Franklin, Javier Abellon-Ruiz, Sophia L. Oram, Lauren Parsons, Alan Cartmell, Gareth S. A. Wright, Arnaud Baslé, Matthias Trost, Bernard Henrissat, Jose Munoz-Munoz, Robert P. Hirt, Laura L. Kiessling, Andrew Lovering, Spencer J. Williams, Elisabeth C. Lowe, Patrick J. Moynihan

## Abstract

Division and degradation of bacterial cell walls requires coordinated action of a myriad of enzymes. This particularly applies to the elaborate cell walls of acid-fast organisms such as *Mycobacterium tuberculosis*, which consist of a multi-layered cell wall that contains an unusual glycan called arabinogalactan. Enzymes that cleave the d-arabinan core of this structure have not previously been identified in any organism. We have interrogated the diverse carbohydrate degrading enzymes expressed by the human gut microbiota and uncovered four families of glycoside hydrolases with the capability to degrade the d-arabinan or d-galactan components of arabinogalactan. Using novel exo-d-galactofuranosidases from gut bacteria we generated enriched d-arabinan and used it to identify *D. gadei* as a D-arabinan degrader. This enabled the discovery of endo- and exo-acting enzymes that cleave D-arabinan. We have identified new members of the DUF2961 family (GH172), and a novel family of glycoside hydrolases (DUF4185) that display endo-d-arabinofuranase activity. The DUF4185 enzymes are conserved in mycobacteria and found in many microbes, suggesting that the ability to degrade mycobacterial glycans plays an important role in the biology of diverse organisms. All mycobacteria encode two conserved endo-d-arabinanases that display different preferences for the d-arabinan-containing cell wall components arabinogalactan and lipoarabinomannan, suggesting they are important for cell wall modification and/or degradation. The discovery of these enzymes will support future studies into the structure and function of the mycobacterial cell wall.

## Introduction

Growth and division of all bacteria is a carefully orchestrated process requiring the coordinated action of a host of enzymes. Acid-fast organisms such as *Mycobacterium tuberculosis* possess unusual cell wall glycans and lipids which require additional enzymatic machinery during this process. The core of the cell wall structure is conserved amongst mycobacteria and consists of three layers^1,2^. Like other bacteria, peptidoglycan forms the basal layer of the cell wall, though the precise architecture is unknown. At the other extremity of the wall are the mycolic acids which give the organisms their characteristic waxy appearance and are interspersed with a host of species-specific free lipids. Joining these two layers is a complex polysaccharide called arabinogalactan (AG), which has a chemical composition unique to the Mycobacteriales and entirely distinct from the similarly named molecule found in plants that is composed of l-arabinofuranose (**Figure 1A**)^3^. AG is comprised of two domains with a β-d-galactofuranose backbone decorated by large α-d-arabinofuranose branches^4^. The structure and biosynthesis of this molecule has undergone intense scrutiny, due in part to it being a target of the antimycobacterial drug ethambutol^5–8^. Ethambutol targets the arabinosyltransferase proteins in the cell envelope of mycobacteria that are responsible for biosynthesis of the polysaccharide^9^. Similarly, the biogenesis of mycolic acids and peptidoglycan are the subject of much research due to their biochemical complexity, essentiality, and their synthesis being the target of isoniazid and β-lactams respectively^10,11^.

**Figure 1.**
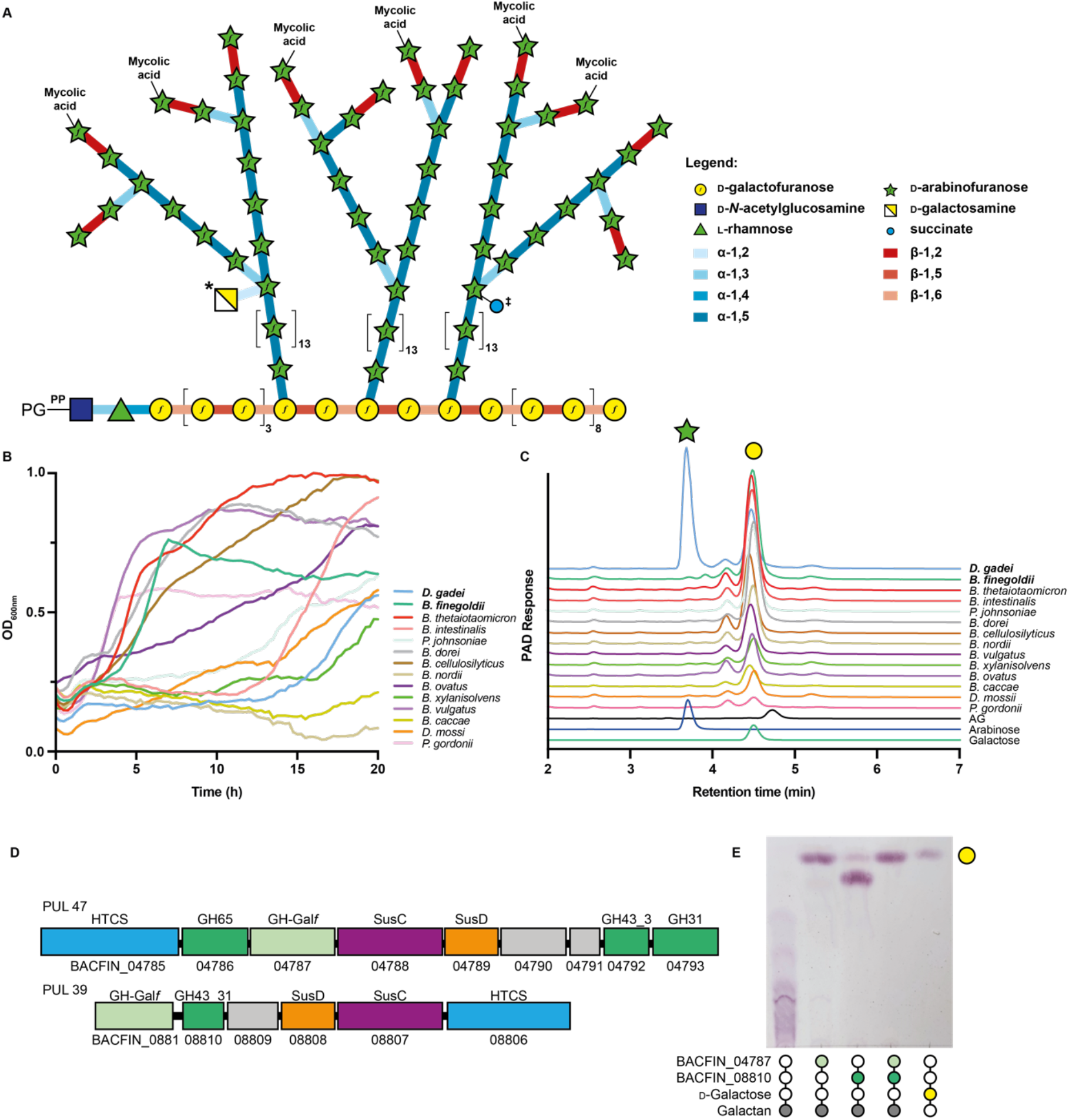
Bacteroidetes growth on mycobacterial arabinogalactan. **A)** Schematic of the structure of mycobacterial mycolyl-arabinogalactan-peptidoglycan complex. Succinate (‡) and galactosamine (*) modifications are non-stoichiometric. **B)** Growth of selected Bacteroidetes species on arabinogalactan as monitored by change in OD_600nm_. **C)** IC-PAD analysis of culture supernatants for bacteria grown on mycobacterial AG as a sole-carbon source. Production of arabinose (green star) and galactose (yellow circle) are indicated and compared to standards. **D)** Schematics of PUL47 and 39 from *B. finegoldii*, whose homologs in *B. cellulosilyticus* were identified by RNAseq as being upregulated during growth on AG. **E)** TLC analysis of endpoint reaction products of *B. finegoldii* enzymes with *C. glutamicum* Δ*ubiA* galactan. Filled circles indicate the presence of a given reaction component.

The growth of mycobacteria likely requires not only the coordinated synthesis of the cell wall, but also its eventual degradation and turn-over to allow for cellular expansion, division and the insertion of cell envelope spanning secretion systems. Similarly, the control of cell surface structures affords these bacteria the ability to modulate how they are sensed by the host. As such, hydrolases of the cell wall may play an important role at the mycobacterial cell surface. Hydrolases of peptidoglycan influence mycobacterial morphology and infection and it was recently demonstrated that at least some peptidoglycan by-products are recycled by the bacteria^12,13^. Similarly, trehalose-mycolates, a dominant lipid component of the cell wall, undergo recycling,^14^ and a hydrolase involved in turning over mycolic acids (Rv3451) and a recycling pathway for trehalose has been described^15^. Cleavage of the acid-fast cell wall is unlikely to be restricted to the Mycobacteriales. Organisms that predate on acid-fast bacteria such as phage or bacterial predators like the recently reported *Candidatus* Mycosynbacter amalyticus (hereafter M. amalyticus), are also likely to require D-arabinan degrading enzymes to penetrate the mycobacterial cell wall^16^.

Only a single enzyme capable of degrading AG has been characterized to date. GlfH1 (Rv3096) is an exo-β-d-galactofuranosidase from glycoside hydrolase (GH) family GH5_13 that cleaves the galactan backbone of mycobacterial AG at both β-1,5 and β-1,6 linked residues^17^. The precise role of this enzyme in mycobacterial biology remains unclear but its activity is suggestive of it being part of a remodeling pathway for this cell wall component. Enzymes with activity on simple d-galactofuranosides (d-Gal*f*) have been identified through large-scale screening of orphan GH sequences^18^. Likewise, no enzymes have been characterized with exo- or endo-D-arabinofuranase activity (enzymes that cut within the D-arabinan polymers) against AG, although this activity was described in protein extracts from soil bacteria dating back to the 1970s^19,20^. Endo-arabinan activity was also described in extracts of *Mycobacterium smegmatis and* was suggested to increase upon treatment of cells with ethambutol, which blocks D-arabinan biosynthesis^21,22^. During our studies an exo-acting difructose-dianhydride I synthase/hydrolase was discovered that was active on *p*NP-α-d-arabinofuranoside^23^. Whether this enzyme acts on mycobacterial AG is unknown.

The human gut microbiota is responsible for the degradation of dietary plant polysaccharides, host and microbial glycans. Dominated by the Bacteroidetes, this grouping of organisms is collectively amongst the richest known organisms in diversity of complex carbohydrate degrading enzymes^24^. Carbohydrate utilization by the Bacteroidetes is typically mediated by genes which are organized into polysaccharide utilization loci (PULs), which can be induced upon exposure of the bacterium to a given carbohydrate^25–28^. The abundance and diversity of carbohydrate degrading enzymes in the Bacteroidetes provides a rich opportunity for enzyme discovery.

In this study we have mined the glycolytic capacity of the human gut microbiota and discovered a collection of enzymes able to completely degrade mycobacterial arabinogalactan. We report the discovery of new families of glycoside hydrolases active on the mycobacterial cell wall. We also identify new exo-d-arabinofuranosidases from the DUF2961 family (GH172). Furthermore, we demonstrate that D-arabinan degradation is wide-spread amongst Mycobacteriales, and mycobacteria but is also present in phages, other bacteria, and microbial eukaryotes. Our data point to a key role for these enzymes in mycobacterial biology and can enable sophisticated analysis of mycobacterial cell wall components.

## Results

### Select human gut Bacteroidetes can utilize mycobacterial AG as a carbon source

While endo-d-arabinofuranase activity was first described in soil bacteria more than 50 years ago and has been known in mycobacteria for at least 30 years^19,20^, the enzymes responsible for this activity have escaped identification. We reasoned that these enzymes might be highly regulated, unstable, or poorly soluble in mycobacteria making purification-based approaches unfeasible. Moreover, if they belong to novel enzyme class(es), bioinformatics approaches would fail. Instead, we isolated arabinogalactan from *M. smegmatis* mc^2^155 and used it as sole-carbon source for the growth of a panel of 14 Bacteroidetes species. Of these, 12 strains were able to grow on this material (**Figure 1B**). Ion chromatography with pulsed amperometric detection (IC-PAD) analysis of selected culture supernatants (**Figure 1C**) demonstrated the production of free galactose in most cultures, and arabinose in one; *Dysgonomonas gadei*. Together these data indicate that members of the gut microbiota produce enzymes that can depolymerize mycobacterial d-arabinan and d-galactan.

### Identification of exo- and endo-galactofuranosidases that degrade galactan

The presence of both galactose and arabinose in *D. gadei* culture supernatants complicated the identification of PULs specific for either galactan or arabinan. Therefore, we developed a method for production of pure D-arabinan by exploiting d-galactan specific PULs. Based on the analysis of culture supernatants, *B. finegoldii* and *B. cellulosilyticus* appeared to degrade galactan, but not d-arabinan. RNAseq analysis of *B. cellulosilyticus* revealed the upregulation of PUL35 and PUL36, containing multiple predicted GHs (**Figure S1**). While we could not heterologously express the *B. cellulosilyticus* enzymes, the homologs of these enzymes from *B. finegoldii* DSM17565 (PULDB ID: PUL39 and PUL47, **Figure 1D**) could be expressed and purified, and when tested on galactan (**Figure 1E**) demonstrated exo- and endo-activities. To determine their galactan degradative capacity, we purified galactan, comprised alternating β-1,5- and β-1,6-galactofuranose residues from a strain of *Corynebacterium glutamicum* (Δ*ubiA*) that lacks d-arabinan^29^. As shown in **Figure 1E** and **Figure S1**, combination of the two *B. finegoldii* enzymes comprising a GH43_31 (BACFIN_08810) and a new GH-exo-Gal*f* (BACFIN_04787) family completely hydrolysed this galactan substrate.

### Cultivation on d-arabinan identifies D-arabinan-active PULs

To generate d-arabinan we digested mycobacterial AG with the two *B. finegoldii* galactofuranosidases (BACFIN_08810 and BACFIN_04787). This resulted in an enriched D-arabinan fraction with approximately 70% reduction in galactan as determined by acid hydrolysis (**Figure S1**). The resulting d-arabinan was used as a sole carbon source for *D. gadei* (which was previously shown to produce both d-galactose and d-arabinose from AG). Proteomics of these bacteria at mid-log phase during growth on enriched D-arabinan identified a predicted fucose isomerase as the most abundant carbohydrate-active enzyme (**Table S1**). This protein maps to PUL42 in the *D. gadei* genome (**Figure 2A**), and an additional nine of the proteins derived from this PUL were in the top 200 most abundant proteins in the total proteome, including several that lacked annotation (**Figure S2**). Reasoning that mycobacteria may harbor homologs of *D. gadei* arabinanases to process arabinan, we prioritized the DUF2961 and DUF4185 superfamily enzymes encoded in PUL42 as they possessed homology to predicted mycobacterial proteins of unknown function within the same superfamilies.

**Figure 2.**
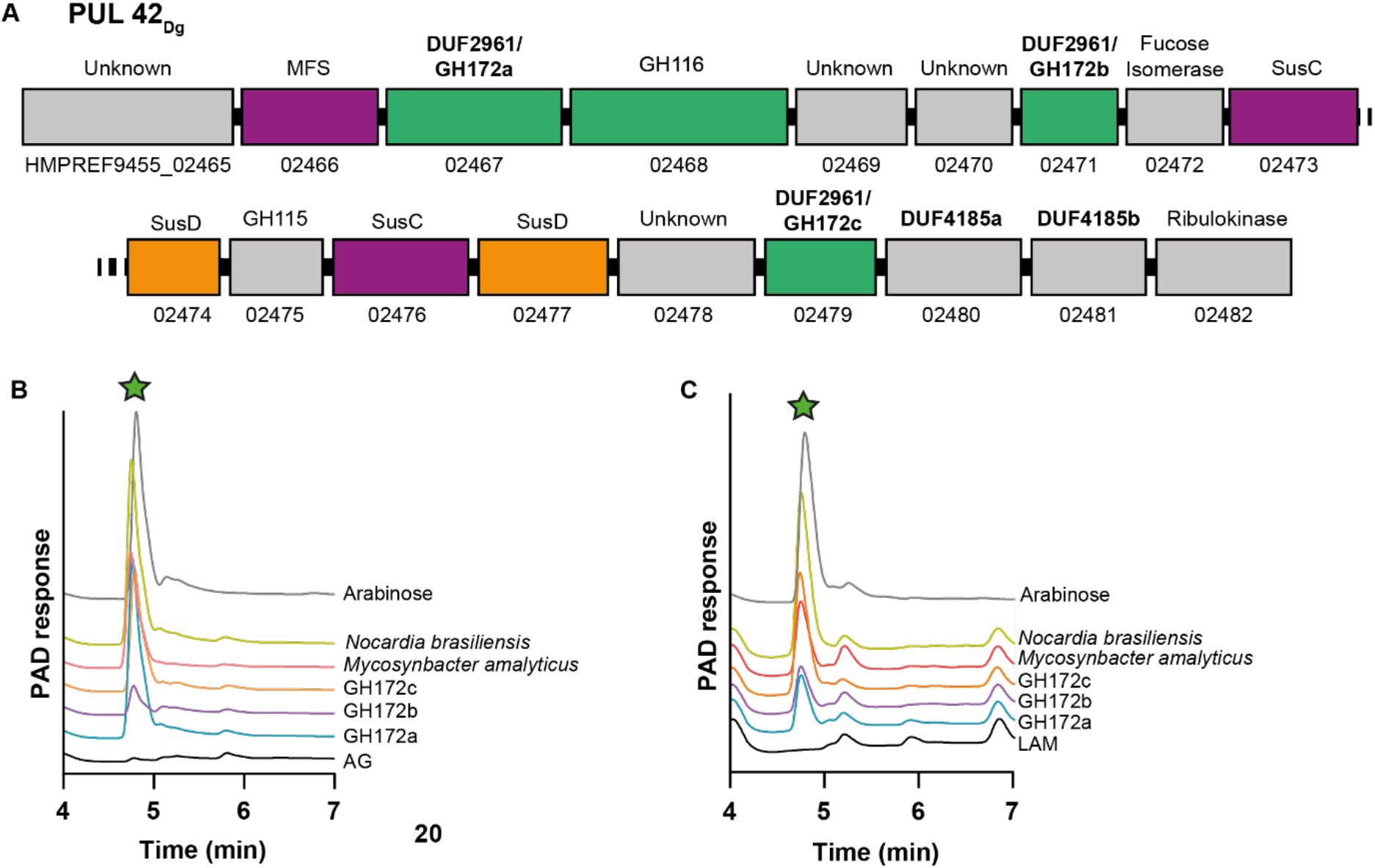
*D. gadei* DUF2961 genes encode GH172 exo-D-arabinofuranosidases with orthologs in diverse bacteria. **A**) Schematic of PUL42 as identified by proteomic analysis of *D. gadei* grown on d-arabinan. **B**) DUF2961 enzymes (1 μM) vs 2 mg ml^−1^ AG and **C**) LAM. Samples were analyzed on a Dionex ICS-6000 with CarboPac Pa300 column, 100 mM NaOH, 20 min isocratic elution followed by a 0-60% 500 mM sodium acetate gradient over 60 minutes. Green star = d-arabinose.

### The DUF2961 superfamily (GH172) includes d-arabinofuranosidases

At the outset of this study, no member of the DUF2961 family had been characterized. Therefore, we cloned, expressed, and purified all three DUF2961 family members found in PUL42 in *D. gadei*. Upon incubation with purified AG, we observed d-arabinofuranosidase activity for HMPREF9455_02467, HMPREF9455_02471 and HMPREF9455_02479 (hereafter Dg_GH172a_, Dg_GH172b_ and Dg_GH172c_, respectively) **(Figure S3)**. To probe the activity of these enzymes, we synthesized the chromogenic substrates *p*-nitrophenyl α-d-arabinofuranoside (pNP-α-d-Ara*f*) and β-d-arabinofuranoside (pNP-β-d-Ara*f*). Dg_GH172a_ and Dg_GH172c_ were active against pNP-α-d-Ara*f*, and no activity was observed using pNP-β-d-Ara*f* (**Table 1**). Although substrate limitations prevented accurate determination of *V*_max_ we could use pNP-α-d-Ara*f* to determine *k*_cat_/*K*_M_ for Dg_GH172c_ (**Figure S4 and Table 2**). These data demonstrate that gut bacteria can use DUF2961 enzymes to generate d-arabinose from mycobacterial arabinogalactan.

**Table 1:** Activity of GH172 enzymes against synthetic and bacterial substrates.

| Protein | pNP-α-D-Araf | AG | LAM | Pili |
| --- | --- | --- | --- | --- |
| **Dg**<sub>GH172a</sub> | ✓ | ✓ | ✓ | ✓ |
| **Dg**<sub>GH172b</sub> | weak | ✓ | ✓ | weak |
| **Dg**<sub>GH172c</sub> | ✓ | ✓ | ✓ | ✓ |
| **Noc**<sub>GH172</sub> | ✓ | ✓ | ✓ | ✓ |
| **Myc**<sub>GH172</sub> | ✓ | ✓ | ✓ | weak |

**Table 2:** *k*_cat_/*K*_M_ of GH172 enzymes for pNP-α-d-Ara*f* and AG.

| Protein | pNP-D-Araf |
| --- | --- |
| **Dg**<sub>GH172a</sub> | n/a |
| **Dg**<sub>GH172c</sub> | $2.9 \times 10^4 \pm 4 \times 10^2 \text{ min}^{-1} \text{ M}^{-1}$ |
| **Noc**<sub>GH172</sub> | $1.2 \times 10^5 \pm 2 \times 10^3 \text{ min}^{-1} \text{ M}^{-1}$ |

### DUF2961 superfamily (GH172) enzymes are present in acid-fast bacteria and their predators

Homologues of the DUF2961 encoding genes are present in the genomes of organisms from the Actinomycetota phylum including pathogens such as *Mycobacterium avium* subsp. *paratuberculosis* (MAP_0339c) and *Nocardia brasiliensis* (O3I_017420; Noc_GH172_). Noc_GH172_ was soluble; however, despite repeated attempts, we were unable to produce soluble MAP_0339c. Noc_GH172_ had d-arabinofuranosidase activity on both pNP-α-d-Ara*f* and purified arabinogalactan (**Table 2, Figure S3**). We also identified a DUF2961 homolog within the genome of the parasitic bacterium M. amalyticus (GII36_05205; Myc_GH172_), which possessed similar activity (**Figure 2 and S3A**)^16^. Together these data indicate that exo-d-arabinofuranosidase activity is a feature of the DUF2961 super-family. These enzymes are encoded by both the Mycobacteriales and their predators.

### GH172 enzymes degrade α-1,5-D-arabinofuranoside linkages within d-arabinan

To understand the substrate specificity of these enzymes against other microbial D-arabinan substrates, we assayed our DUF2961 GH172 enzymes (Dg_GH172a_, Dg_GH172b_, Dg_GH172c_, Noc_GH172_, and Myc_GH172_) against mycobacterial LAM and d-araF containing pilin oligosaccharides (**Table 1**). The d-arabinan branches of lipoarabinomannan (LAM) are structurally conserved with those in AG. All enzymes could digest LAM from *M. smegmatis* to produce arabinose, although Dg_GH172b_ and Myc_GH172_ displayed lower activity than the others (**Figure 2C and Figure S3B**).

Pili from *Pseudomonas aeruginosa* PA7 are decorated with α-1,5-d-arabinofuranose oligosaccharides. All our GH172, except for Dg_GH172b_ enzymes could cleave the d-arabinofuranose oligosaccharides from digested pili (**Figure S3C**). To rule out d-galactofuranosidase activity, we repeated these experiments using purified d-galactan (**Figure S3D**) and observed no activity. The combination of these experiments reveals that GH172 enzymes are specific for α-d-arabinofuranoside linkages, are active on d-arabinan, and can cleave α-1,5-d-arabinofuranoside linkages.

### Family GH172 proteins can adopt diverse oligomeric states

To better understand the structure-function relationship of this protein family, we examined the multimeric state of several of our candidates of interest. Size-exclusion chromatography with laser scattering (SEC-LS) analysis of the GH172 family enzymes allowed assessment of molecular weight and assignment of oligomerization states (**Table 3 and Figure S7**). A wide range of oligomeric states were uncovered: Dg_GH172c_ was assigned a hexamer, Dg_GH172a_ a dodecamer, Noc_GH172_ and Myc_GH172_ as trimers, and Dg_GH172b_ as a dimer. This solution data complements recent reports of hexameric assemblies in protein crystals of two GH172 members: difructose dianhydride I synthase/hydrolase from *Bifidobacterium dentium* (PDB: 7V1V) and BACUNI_00161 from *Bacteroides uniformis* (PDB: 4KQ7) (**Figure S5**)^23^.

**Table 3:**
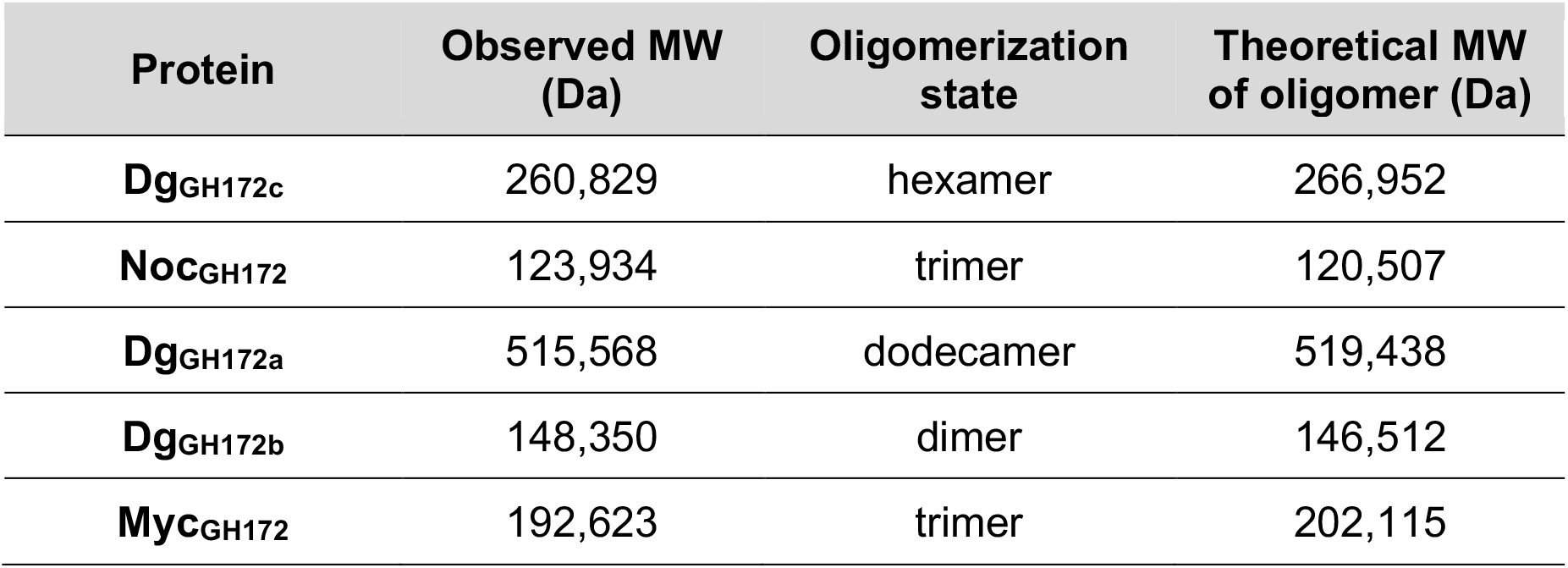
SEC-LS oligomerization state of GH172 enzymes.

Interestingly, Dg_GH172b_ is predicted to have 1.5 DUF2961 domains, and so dimerization of Dg_GH172b_ is predicted to provide three DUF2961 domains in the final assembly. SEC-LS analysis is supportive of a quaternary structure for each of the GH172 proteins exhibiting multiple DUF2961 domains: Dg_GH172b_, Myc_GH172_ and Noc_GH172_ each contain three DUF2961 domains, while Dg_GH172c_ contains six and Dg_GH172a_ contains twelve. Diversity in quaternary structure may lead to differences in activity or substrate specificity.

### The 3D structure of Dg_GH172c_ reveals a dimer-of-trimers assembly

The 3D structure of Dg_GH172c_ was determined by X-ray crystallography using molecular replacement with the *B. uniformis* GH172 (Bu_GH172_, PDB: 4KQ7; 64% sequence identity with Dg_GH172c_) as the search model. The structure of Dg_GH172c_ was refined to 1.4 Å resolution, data collection and model refinement statistics are shown in **Table S2**. The Dg_GH172c_ monomer is comprised of two β jelly rolls and a C-terminal alpha helix (**Figure 3A**). Six monomers assemble in a dimer-of-trimers (**Figure 3B and Figure S5A**), consistent with the solution phase oligomerization state determined by SEC-LS (**Table 3**). The dimer-of-trimers structure is formed by the C-terminal of each monomer of one trimer interlocking with the C-terminal of a monomer from a second trimer. The lower side of the trimers face each other in the dimer-of-trimers structures. The interfaces within the trimer form the active site with residues from both monomers contributing to the active site (**Figure 3C**). This gives a total of 3 active sites per trimer, and six per hexameric assembly. Seven calcium ions are coordinated within each Dg_GH172c_ trimer: two within each active site/protomer, and one in the 3-fold axes.

**Figure 3:**
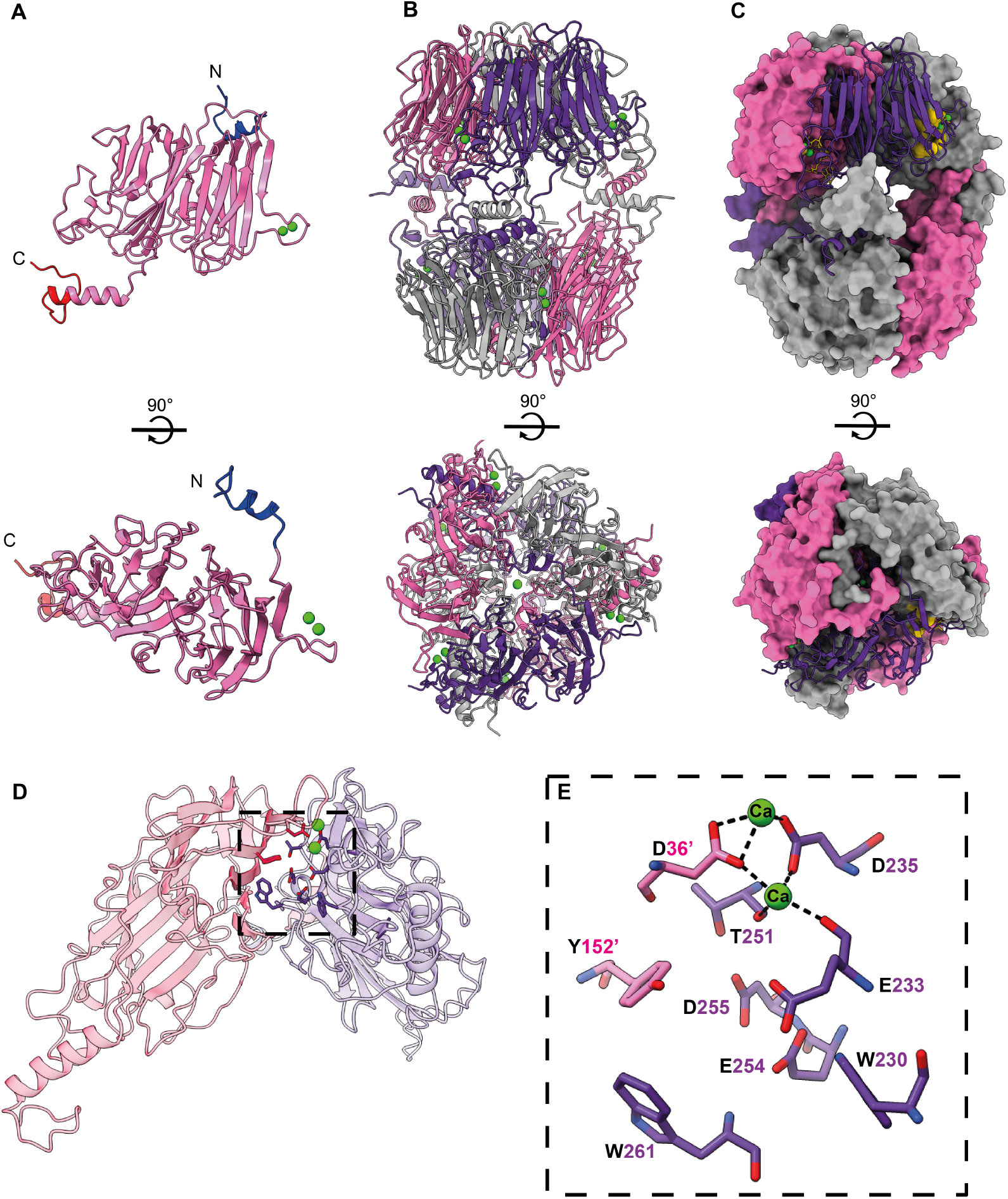
3D structure of Dg_GH172c_ determined by X-ray crystallography. **A)** Monomer of Dg_GH172a_ with secondary structure shown as cartoon. The N- and C-termini are labelled and colored blue and red respectively. Bound calcium ions are shown as green spheres. **B)** Overview of hexameric architecture of Dg_GH172c_. Cartoon representation with bound calcium ions shown as green sphere. Trimer subunits are colored grey, pink and purple for clarity. **C)** Surface representation of the hexamer with one monomer shown in cartoon. Active site residues are colored gold, and calcium ions shown in green. **D)** Cartoon representation of two monomers of Dg_GH172c_ which form the active site, with one monomer in pink and the other in purple. The active site residues and coordinated calcium ions are highlighted by a black dashed box. **E)** Proposed active site residues of Dg_GH172c_. Residues are shown as sticks with carbon atoms colored pink and purple for different subunits. Calcium ions are shown as green spheres and coordinated by D36’, D235 and T251.

The proposed active site of Dg_GH172c_ is formed of residues D36, Y152, W230, E233, D235, T251, E254, D255, and W261 (**Figure 3D**). The proposed catalytic residues identified in the characterization of Bd_GH172_ are conserved in Dg_GH172c_, (Bd/Dg: E270/254, catalytic nucleophile; E291/233, acid/base) and other members of the GH172 family (**Figure S6, S8**). E233 and E254 are 5.4 Å apart in Dg_GH172c_, which is consistent with a retaining mechanism of enzyme catalysis, as shown for Bd_GH172_^23^. Ca^2+^ ions are observed in the active sites of Dg_GH172c_ (contains two Ca^2+^) and Bd_GH172_ (contains one Ca^2+^). The calcium ions in Dg_GH172c_ are coordinated by D235, which is not conserved in Bd_GH172_. Instead, at the equivalent position, Bd_GH172_ has N272. A particular feature of characterized GH172 family members is the formation of the active site by two monomers, an uncommon feature of GHs, but on that has been observed in *E. coli* LacZ β-galactosidase^30^.

### Dg_GH172c_ catalysis is driven by conserved glutamate residues

To gain insight into the functional roles of the residues in the active site of DgGH172c, the proposed catalytic carboxylate residues were mutagenized to alanine, generating variants E233A, E254A and D225A. Dg_GH172c_-E233A and Dg_GH172c_-D225A had no detectable activity and there was a greater than 10^5^-fold reduction in activity for Dg_GH172c_-E254A. Our data support the assignment of E254 and E233 as catalytic residues (acid/base or nucleophile) and highlight an important role for the adjacent D255 residue. None of the variants displayed a change in oligomerization state suggesting they do not possess structural roles (**Figure S9**).

### DUF4185 is widely distributed throughout bacterial species

In Bacteroidetes, the degradation of a target polysaccharide is typically a multi-step process whereby oligosaccharides are generated and subsequently cleaved into their monosaccharide constituents, encoded within co-transcribed operons. We reasoned that the generation of D-arabinofuranose oligosaccharides was likely achieved by proteins that have no annotated function and so focused our attention on the DUF4185 proteins. To investigate the conservation of these genes across different organisms, we constructed a phylogeny of DUF4185 proteins. This revealed that they are common within actinobacteria, especially amongst the Actinomycetota (**Figure 4 and Figure S10**), in *Bacteroides* species, *Myxococcus* and in the lysis cassette of some actinobacteriophage (**Figure S11**) and the predacious bacterium M. amalyticus.

**Figure 4:**
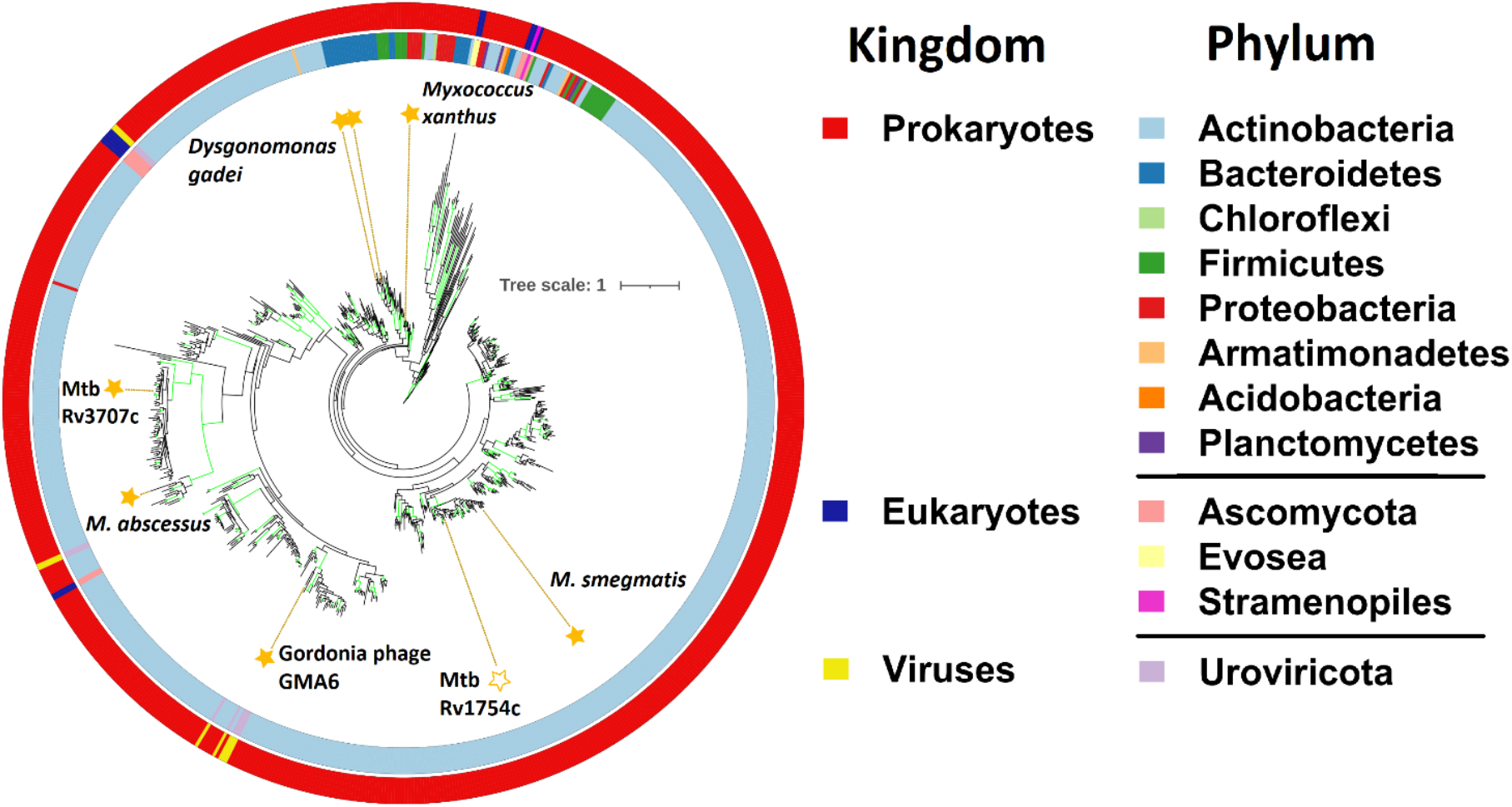
Phylogeny of the DUF4185 enzyme family. Unrooted ML phylogeny (LG model with empirical base frequencies, invariable sites and the discrete gamma model) of DUF4185 family sequences. Branches with greater than 75% bootstrap support (100 replicates) are highlighted in green. Units for tree scale are inferred substitutions per amino acid residue. Colored rings indicate phylum (inner) and kingdom (outer) taxonomy information for sequences. Stars highlight sequences of interest and are filled for proteins that have been experimentally characterized in this work.

### DUF4185 comprises a novel GH family with endo-d-arabinanase activity

Initially, we cloned, expressed, and purified the DUF4185 homologs (HMPREF9455_02480 and HMPREF9455_02481) from *D. gadei* (herein, referred to as Dg_GH4185a_ and Dg_GH4185b_, respectively). When incubated with mycobacterial AG and then analyzed by HPAEC, these enzymes produced a banding pattern characteristic of an endo-acting GH (**Figure S12A**), consistent with endoarabinanase activity. To assess the breadth of activity for DUF4185 enzymes we selected additional candidate enzymes from each of the major lobes of the global phylogeny (**Figure 4**). Recombinant proteins were produced from several bacterial lineages and a phage capable of infecting Gram-positive bacteria *Gordonia* of the order Mycobacteriales. These included *Myxococcus xanthus* (Myxo_GH4185_), M. amalyticus, and *Gordonia* Phage GMA6 (Phage_GH4185_). Where the DUF4185 domain sat within a larger gene containing several other large domains we produced truncated variants containing only the DUF4185 domain due to low solubility of the multi-domain proteins. All DUF4185 constructs except that from M. amalyticus yielded soluble protein. As shown in **Figure 5** and **Figure S12**, when incubated with mycobacterial AG all these enzymes possessed endo-d-arabinanase activity with varying product profiles suggesting differences in enzyme specificity. Some of the enzymes were also active against the linear α-1,5-d-arabinofuranose oligosaccharides derived from *P. aeruginosa* PA7 pili, consistent with activity against the major linkage of mycobacterial d-arabinan (**Figure S12**). The discovery of endo-d-arabinanase activity in proteins from organisms outside of the Actinomycetota is an interesting observation, however its presence in bacterial predators is consistent with the essentiality of this polymer for mycobacterial viability. This sugar motif has been reported in a small number of LPS structures, providing a possible explanation for the presence of this enzymatic activity in the gut microbiota^31,32^. Furthermore, some corynebacterial species are found as commensals of the human oral and gut microbiome^33^.

**Figure 5:**
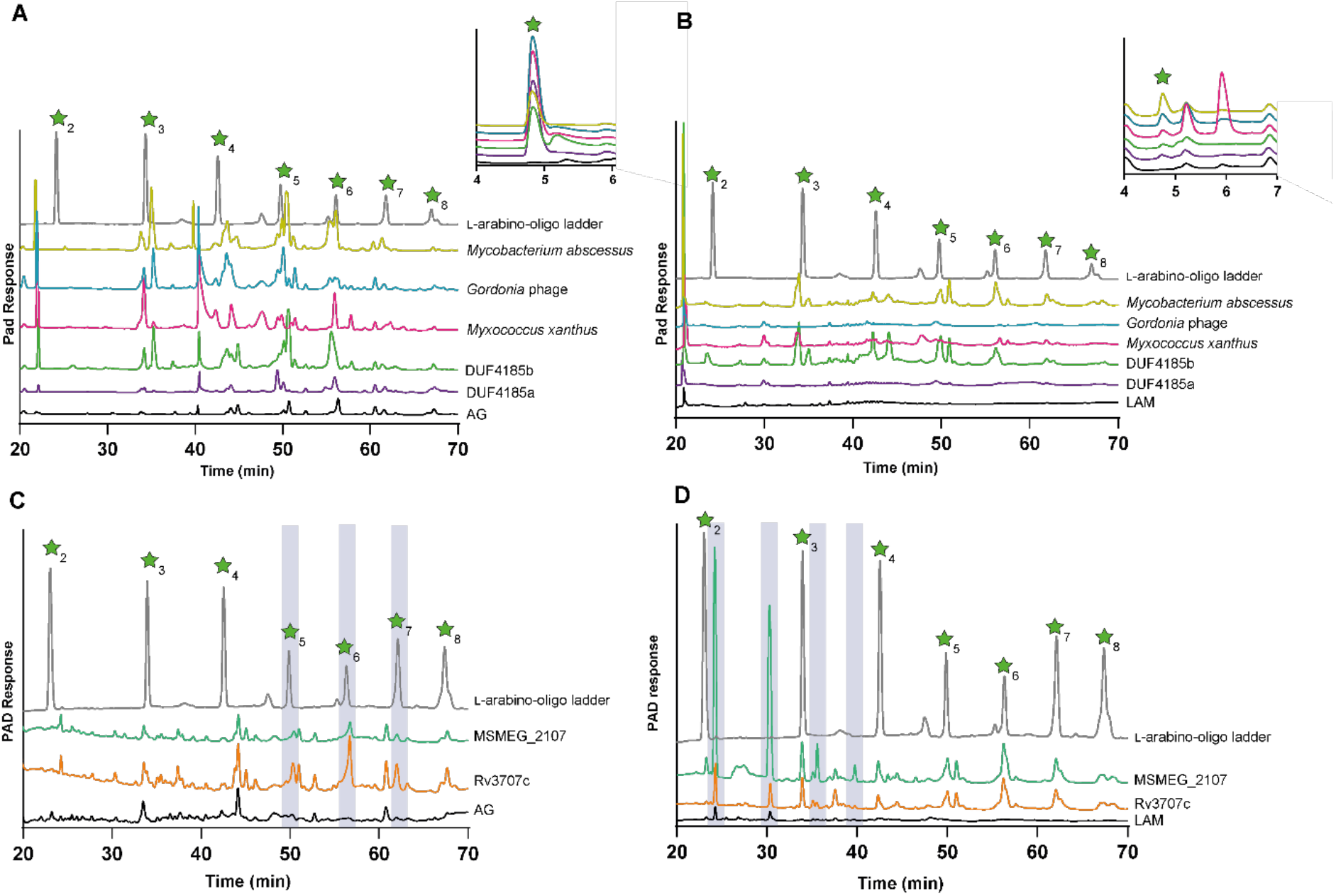
DUF4185 proteins are endo-D-arabinofuranases that cleave mycobacterial AG and LAM. DUF4185 enzymes were incubated with 2 mg·ml^−1^ AG (**A** and **C**) or 2 mg· ml^−1^ LAM (**B** and **D**) for 16 hours as described in Materials and Methods. Samples were analyzed by High-performance anion-exchange chromatography/pulsed amperometric detection (HPAEC-PAD) on a Dionex ICS-6000w with CarboPac Pa300 column, 100 mM NaOH 20 min isocratic elution followed by a 0-60% 500 mM sodium acetate gradient over 60 minutes. A ladder of α-1,5-l-arabino-oligosaccharides (25 μM) derived from plant arabinogalactans was used as a standard. In panel C the chromatogram of this ladder has been scaled on the *y*-axis by a factor of 0.2 for clarity of presentation.

### Mycobacterial DUF4185s are endo-d-arabinofuranases

Given the importance of d-arabinan to mycobacterial viability and immunology we next sought to understand the biochemical function of DUF4185 proteins from representative mycobacteria. As shown in **Figure 4** and **Figure S10**, mycobacteria produce at least two DUF4185s that fall into distinct phylogenetic groupings. In *M. tuberculosis* these are Rv1754c and Rv3707c. Beyond these two conserved DUF4185 genes, some species additional DUF4185 members. For example, many *Mycobacterium abscessus* strains encode at least three distinct members whilst *M. smegmatis* mc^2^155 encodes five (MSMEG_4352, 4360, 4365, 2107 and 6255). Based on sequence analysis, MSMEG_2107 and MSMEG_6255 are homologs of Rv1754c and Rv3707c, respectively, while the remainder show greater diversity (**Figure S10**). A distant homolog of Rv3707c from *Mycobacterium abscessus*, Ga0069448_1118 (hereafter referred to as *Mab*_4185_), was readily produced in soluble form and in good yield^34^. Despite low sequence identity (33.5%) (Figure S13) to the *D. gadei* enzymes, the TLC and HPAEC profiles of mycobacterial AG digested by Mab4185 indicate that it also possesses endo-d-arabinofuranase activity (**Figure 5**).

Encouraged by this success with *Mab*_4185_ but recognizing it has limited sequence identity with either of the *M. tuberculosis* proteins (17% and 16.4% identical to Rv3707c and Rv1754c respectively over the entire protein length), we sought to study Rv3707c and Rv1754. However, despite our best efforts we were unable to produce usable amounts of soluble Rv3707c and Rv1754. A previous report highlighted that Rv3707c is secreted, but it lacks a discernible signal peptide^35^. We reasoned that the instability of Rv3707c may be due to incorrect annotation of the start site in the *M. tuberculosis* H37Rv genome, given that mycobacteria frequently use TTG and GTG start codons in addition to the canonical ATG. We re-evaluated the genomic context of the protein and identified several possible in-frame N-terminal extensions of the gene (**Figure S14**). We compared these potential N-terminal extensions to alignments of DUF4185 proteins that possessed a second, N-terminal domain to identify sequence motifs that are conserved; in many cases a proline-rich region was observed, which is also found in the potential N-terminal extensions of Rv3707c (**Figure S14**). Furthermore, by extending the N-terminus of the protein, a putative signal peptide can be predicted by SignalP 6.0 (**Figure S14**). While the precise start-site is uncertain, the likely cleavage point for the signal peptide could be confidently assigned. Therefore, we designed an expression construct with the putative signal peptide removed, but the remainder of the proline-rich N-terminus intact. This produced a reasonably stable and soluble Rv3707c protein in good yield that digested AG to give products consistent with a hepta- or hexa-saccharide, and was also active on LAM (**Figure 5C/D**). While we were unable to produce soluble Rv1754c, we successfully produced the *M. smegmatis* homolog (MSMEG_2107), which demonstrated activity against LAM, but not AG (**Figure 5C/D and Figure S12**).

### DUF4185 family members from different clades have distinct substrate specificities

To further probe the substrate specificity of these enzymes and elucidate the function of MSMEG_2107, we incubated each of the DUF4185 enzymes (Dg_GH4185a_, Dg_GH4185b_, Mab_GH4185_, Phage_GH4185_, Myxo_GH4185_, MSMEG_2107, and Rv3707c) with: LAM (**Figure S12B**); purified d-galactan (**Figure S12C**); and pilin oligosaccharides from *P. aeruginosa* PA7 (**Figure S12D**). The majority of the enzymes displayed higher activity against AG than LAM. Conversely, MSMEG_2107 was unique in displaying a preference for LAM with no detectable activity when incubated with AG under similar conditions. To confirm that the enzymes only degraded d-arabinan and not d-galactan we tested their activity with d-galactan but did not observe any product formation (**Figure S12C**). Given that branched d-arabinan is the only conserved structure within both AG and LAM, we conclude the DUF4185 family are endo-d-arabinofuranases. We note that only a subset of the DUF4185 enzymes were active against pili from *P. aeruginosa* PA7. As these oligosaccharides are relatively short and linear, this suggests specific enzyme subsite occupancy for activity, which leads to production of arabinose for some enzymes; by contrast arabinose production was not observed upon digestion of branched, polymeric AG.

We next assessed whether DUF4185 endo-d-arabinofuranases can cleave AG in the context of an intact cell wall. We utilized metabolic arabinogalactan labelling with the azide-modified lipid-linked Ara*f* donor 5-AzFPA and then fluorescently labelled intact bacteria with DBCO-conjugated AF647 using click chemistry (**Figure S16**)^36–38^. Both Mab_GH4185_ and Rv3707c released fluorescently labelled material from cell wall, with greater activity and a greater range of products formed by the former enzyme, as observed by HPAEC-PAD. Likewise, treatment with GH172_*Noc*_ led to release of fluorescent products, supporting the conclusion that both groups of enzymes can cleave AG within the mycobacterial cell wall.

### Rv3707c has a highly flexible structure and belongs to the beta-propeller superfamily of glycoside hydrolases

Attempts were made to crystallize all the DUF4185 homologs assayed above. *M. tuberculosis* Rv3707c yielded crystals that diffracted to a resolution of 2.4 Å. Using the AlphaFold structural prediction as a search model for molecular replacement, we solved a partial structure. As shown in **Figure 6A**, Rv3707c possesses a beta-propeller structure, similar to several other glycoside hydrolase families,^39^ with each blade consisting of 3 antiparallel beta sheets organized radially around a central pore. Poor or no density was obtained for amino acid residues 1-9, 22-38, 60-69, 85-86, 285-300, and 325-337, which are located on loops surrounding the predicted active site. The lack of density suggests significant structural flexibility in these regions, consistent with variations in AlphaFold predictions for both Rv3707c and Rv1754c (**Figure 6, S15, S17**).

**Figure 6:**
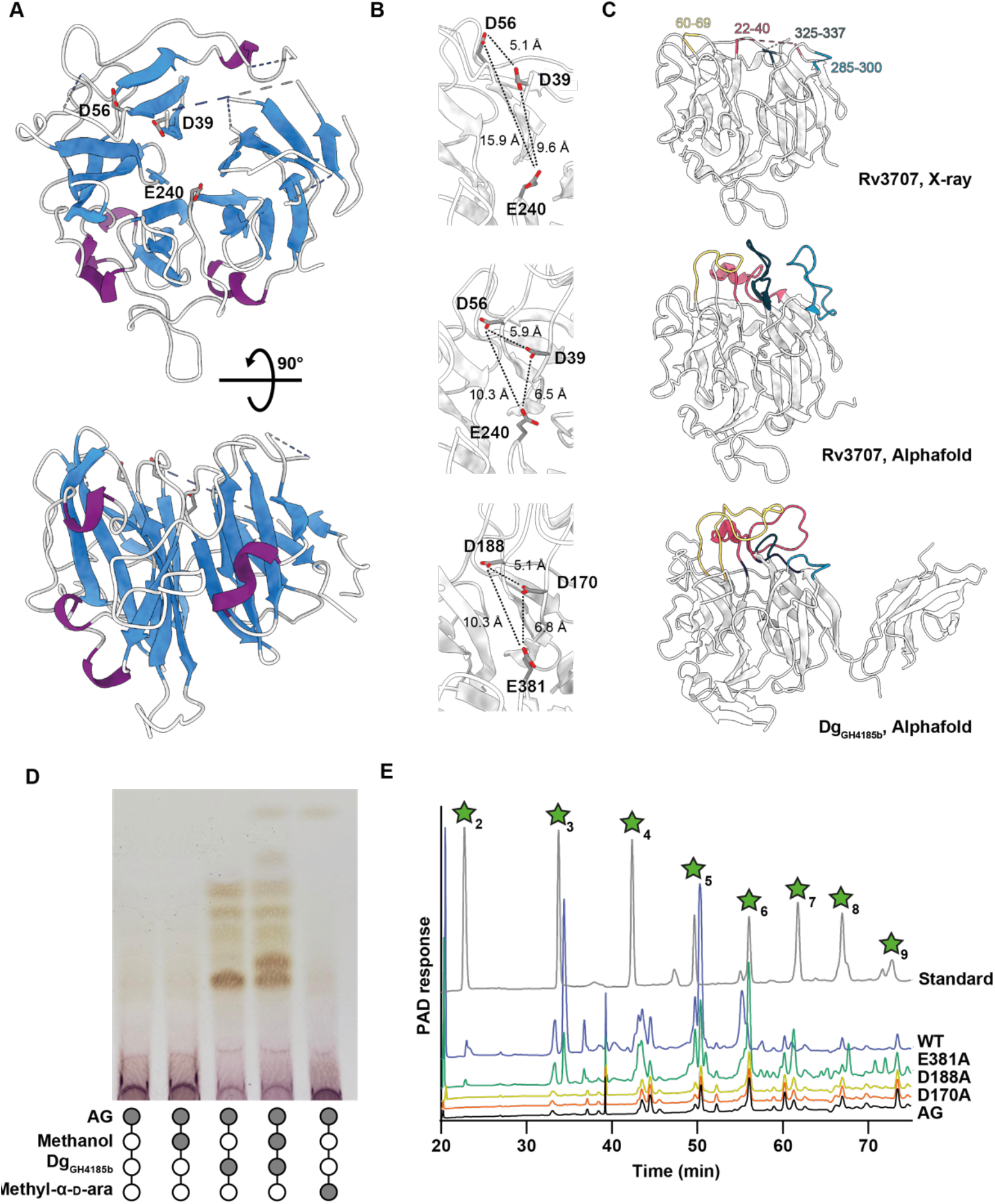
3D structure, active site, and AG digestion by DUF4185 endo-D-arabinofuranases. **A)** 3D structure of Rv3707c determined by X-ray crystallography has a 5-bladed beta-propeller fold. Conserved acidic residues D39, D56 and E240 are indicated in the proposed active site. **B**) Active site details for Rv3707c (X-ray), Rv3707c (AlphaFold) and Dg_GH4185b_ (AlphaFold) with distances between residues indicated in angstroms. **C**) Overall fold of the same enzymes as in **B** with variable surface features highlighted relative to the Rv3707c X-ray structure. Corresponding regions are in the same color in each protein. **D)** Methanolysis of AG catalyzed by Dg_GH4185b_. The production of methyl glucosides in the presence of methanol indicates a retaining mechanism. **E)** Dg_GH4185b_-WT, Dg_GH4185b_-D149A and Dg_GH4185b_-D167A were incubated at 1 μM with 2 mg ml^−1^ AG for 16 hours. No activity was observed for the D188A or D170A mutants.

Multi-sequence alignments of DUF4185 homologs (**Figure S13**) identified three nearly invariant acidic residues in the active site: Asp39, Asp56, and Glu240 (**Figure 6B**). Comparing our experimental structure to the AlphaFold prediction model, the loops that contained the conserved aspartate residues in the proposed active site deviate in their positions (**Figure 6B and Figure S17**). The conserved D56 points away from the active site in our experimental structure, likely in a catalytically incompetent position, while the D39 lies in a similar, but distinct position to that of the AlphaFold model. Together, these observations suggest that the active site may rearrange upon binding of substrate.

The other regions of incomplete density in the experimentally determined 3D structure include loops that are predicted by AlphaFold to be flexible, which is reflected in the B-factors for the experimental structure (**Figure S17**). These regions are also predicted by AlphaFold to be variable in other DUF4185 proteins, including Dg_GH4185b_ (**Figure 6C**). This apparent flexibility likely enables adaptive binding to a range of substructures within highly branched mycobacterial d-arabinan, supporting the production of a broad reaction product profile, as seen in enzymatic digests (**Figure 5**).

### DUF4185 family are anomer-retaining enzymes

Glycoside hydrolases can hydrolyze the anomeric linkage through either reversion or retention. Inclusion of a simple alcohol, such as methanol, in an enzymatic digest can be used to identify retaining enzymes, as they may afford methylated glycosides^40^. Addition of methanol to AG digests by Dg_GH4185b_ afforded methyl arabinoside, thereby demonstrating a retaining mechanism (**Figure 6D**). Three conserved carboxylate residues are present in the active site (**Figure 6B**). Using site-directed mutagenesis, we varied these carboxylate residues in Dg_GH4185b_ to generate Dg_GH4185b_-D170A and Dg_GH4185b_-D188A and Dg_GH4185b_-E381A derivatives. Activity was broadly retained for glutamate substitution, but no activity was observed for the two aspartate mutants (**Figure 6E**). These data support a retaining mechanism for the DUF4185 family of enzymes and assignment of D170/D39 in Dg_GH4185b_ (and D188/D56 in Rv3707c) as the catalytic residues corresponding to acid/base and nucleophile.

## Discussion

Endo-d-arabinanase activity was first reported more than 50 years ago, but despite the widespread availability of mycobacterial genomic tools and -omics technologies, these enzymes have escaped identification^22,41^. We have mined the human gut microbiome and leveraged evolutionary conservation to identify these enzymes in mycobacteria, along with new exo-d-arabinofuranosidases and exo-d-galactofuranosidases (**Figure 7**). We reasoned that the abundance of Mycobacteriales in environmental niches in addition to the availability of d-arabinofuranose polymers in organisms such as *P. aeruginosa* PA7 and corynebacteria meant that the capacity to degrade this carbohydrate was likely to be encoded in the human gut microbiota^42,43^, allowing identification by metabolic methods with arabinogalactan as sole carbon source.

**Figure 7.**
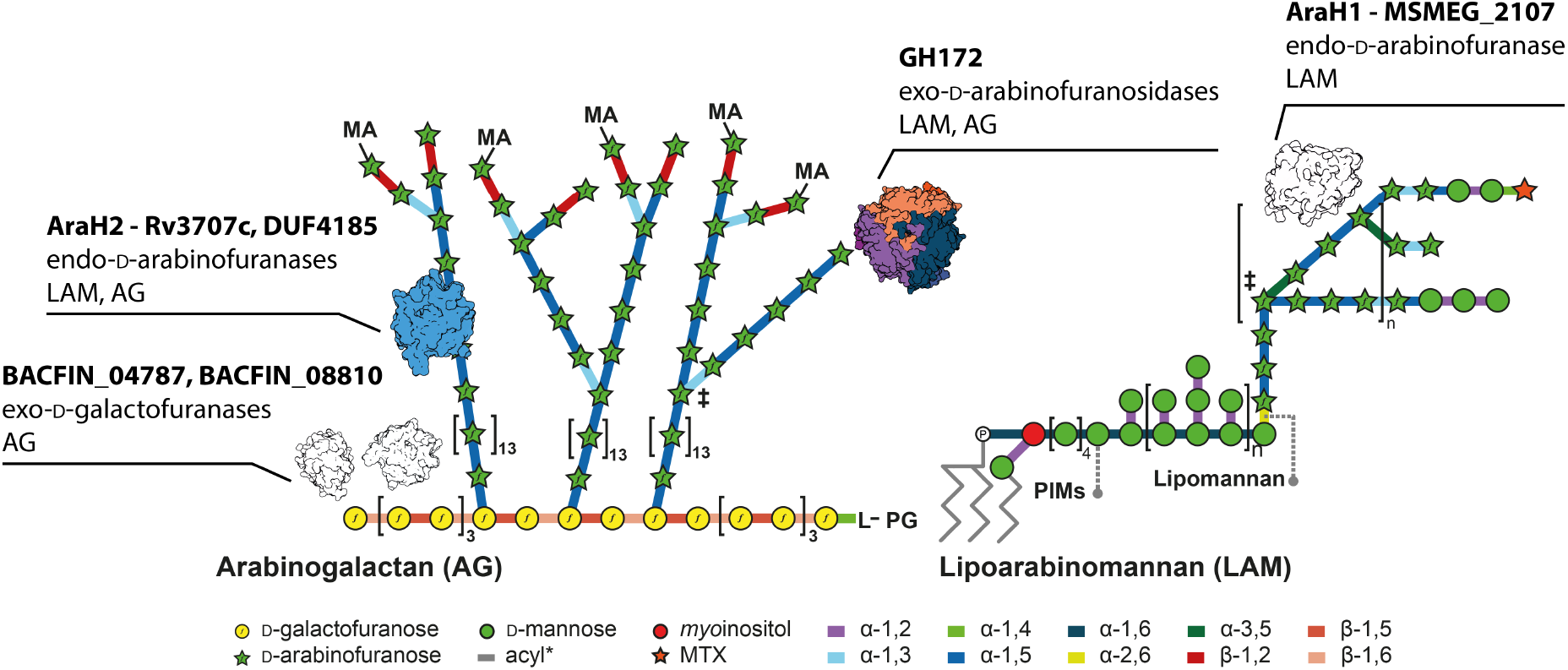
Mycobacterial arabinogalactan-degrading enzymes discovered in this study and their substrates. Each enzyme or enzyme family is listed with its identified function and the mycobacterial cell wall it acts upon. Colored in structures are those for which we have presented experimental structural data, black and white are surface representations of AlphaFold models. MA – mycolic acids; L – linker unit; PG – peptidoglycan; PIMs – phosphatidylinostol mannosides.

Initially, we identified d-galactofuranase activity in a gut microbe organism, whose presence may reflect the widespread occurrence of d-Gal*f* in the LPS of some gram-negative bacteria of the human gut microflora^44^. One of the enzymes we identified, BACFIN_04787, is the founding member of a new glycoside hydrolase family. The galactan degradation pattern exhibited by BACFIN_04787 is consistent with an exo-acting, non-specific enzyme and the product pattern of BACFIN_08810 is consistent with an exo-acting enzyme that can cleave either the β-1,5 or β-1,6 linkage within D-galactan. Further characterization of these enzymes will enable detailed analysis of mycobacterial galactan, whose chain length was recently shown to be important in the biology of these organisms^4^.

Identifying organisms that can grow on d-arabinan is complicated by the presence of d-galactan in AG. To overcome this, we used the newly discovered d-galactofuranase to generate enriched d-arabinan, which we used to identify the Bacteroides *D. gadei*, as a potent d-arabinan degrader. The PULs associated with this ability contained uncharacterized DUFs as well as DU2961 genes corresponding to family GH172. The GH172 proteins were shown to encode exo-d-arabinosidases. During the preparation of this manuscript, members of family GH172 were independently shown to exhibit exo-d-arabinofuranosidase activity by Kashima *et al.*, consistent with our data^23^. The GH172 enzyme from *D. gadei* shares a similar structure to the enzyme described by Kashima *et al*; however, it lacks the large C-terminal extensions observed in Bd_GH172_ which were postulated by the authors to hold the hexameric protein together. SEC-LS analysis of a range of family GH172 proteins reveals diverse quaternary structures of multimers up to dodecamers. How this structural diversity is connected to function is as-of-yet unexplored. It has been suggested that family GH172 proteins and phage capsid proteins may be ancestrally related, raising the possibility for diversification of function within this family^45^.

The presence of genes encoding DUF4185 and GH172 enzymes in M. amalyticus suggests that they contribute to its epibiotic lifestyle on host *Gordonia* spp. through feeding on arabinogalactan as a carbon source. The Myc_DUF4185_ enzyme is predicted to be a surface located lipoprotein, while Myc_GH172_ is predicted to not be secreted. This localization is consistent with a model whereby M. amalyticus releases oligosaccharides from the surface of *Gordonia*, internalizes them, and then cleaves them to monosaccharides in the cytoplasm. An alternative explanation is that the DUF4185 and GH172 enzymes locally remodel the cell wall of *Gordonia*, enabling access to cellular contents. The biological role of GH172 enzymes in *Bacteroides* remains unclear. While some may be involved in α-fructan degradation, those associated with d-arabinan PULs are more likely targeted at either glycans derived from Actinomycetota or organisms such as *P. aeruginosa* PA7.

DUF4185 family of enzymes include the first known endo-D-arabinofuranases, though it is not the first time this activity has been reported. In 1972 Kotani and colleagues reported the isolation of “mixed d-arabinanase activity” in an extract from an unnamed Gram-positive soil microbe, referred to as the “M-2” fraction^19^. This extract possessed endo-activity and released a wide range of products from mycobacterial cell walls, although the specific protein responsible was not identified. Since then, similar impure enzyme cocktails have contributed to numerous studies on AG and LAM^41^. We have now identified broadly conserved endo-d-arabinofuranases from the Mycobacteriales, which we propose to re-name to AraH1 (Rv1754c) and AraH2 (Rv3707c). The availability of well-characterized enzymes with defined activities should support more detailed studies of mycobacterial cell wall polymers.

A review of functional screens of knockout libraries of *M. tuberculosis* highlights an important role for AraH2 in pathogenesis. AraH2 was identified in a screen for proteins with non-canonical signal sequences^35^, likely because of mis-annotation of its signal peptide as demonstrated by our work. In that study, Perkowski and colleagues reported that a transposon mutant in AraH2 was severely defective for replication in macrophages. A separate study identified AraH2 as important for control of phagosome acidification, where it was the second most enriched mutant in an acidified phagosome screen^46^. A homologue of AraH2, PEG_1752, was shown to be upregulated in the phylogenetically related *Mycobacterium llatzerense* upon infection of the amoeba *Acanthamoeba castellanii*^47^. These data point to a role for AraH2 in phagosome survival, and by extension of d-arabinan remodeling in mycobacterial pathogenesis.

The role of AraH1 remains more elusive. The gene is broadly conserved amongst mycobacteria, and our biochemical data suggests this enzyme class is active against LAM, but not AG. This genomic locus is a frequent site of IS6110 element insertion^48^, and interruption of this gene could cause LAM structural variability amongst circulating strains of *M. tuberculosis*. It is possible that AraH1 and AraH2 have partially overlapping substrate specificity and the former can partially compensate for the loss of the latter. As well, these proteins may have specific roles under narrowly defined conditions. Analogously, peptidoglycan-lytic enzymes with seemingly redundant reaction specificities are encoded in most bacteria, and lack notable phenotypes for their loss under most growth conditions. However, screens at a range of pH values have identified specific functions for these enzymes^49^.

To conclude, we have unearthed a new enzymatic toolkit for the degradation of mycobacterial arabinogalactan that should find utility in the study of the structure and function of this important polymer (**Figure 7**). The functional annotation of these genes will support future investigations of the role of d-arabinan structural modulation in mycobacterial biology, mycobacteriophage infection and interbacterial predation.

## Methods

### Bacterial strains and growth conditions

Bacteroidetes sp. were grown on a 2× defined minimal media (**Table S3**) under anaerobic conditions at 37 °C over 24 hours to assay growth on various carbon sources, including arabinogalactan. Strains used were *Bacteroides caccae* ATCC 43185, *B. cellulosilyticus* DSM 14838, *B. dorei* DSM 17855, *B. finegoldii* DSM 17565, *B. intestinalis* DSM 17393, *B. nordii* CL02T12C05, *B. ovatus* ATCC 8483, *B. thetaiotaomicron* VPI-5482, *B. vulgatus* ATCC 8482, *B. xylanisolvens* XB1A, *Dysgonomonas gadei* ATCC BAA-286, *D. mossii* DSM 22836, *Parabacteroides gordonii* DSM 23371, *P. johnsonii* DSM 18315. *Escherichia coli* and *Pseudomonas aeruginosa* PA7 ATCC 15692 strains were grown in lysogeny broth at 37 °C (unless otherwise specified). *Mycobacterium smegmatis* mc^2^155 ATCC 19420 was grown in Tryptic soy Broth at 37 °C with agitation.

### RNA sequencing

*B. cellulosilyticus* was cultured in defined media (supplementary table 3) containing 5 mg ml^−1^ AG or glucose, in triplicate 5 ml cultures. Cells were harvested at mid-log phase and stored in RNA protect (Qiagen). RNA was purified with the RNAeasy Kit. Prior to library preparation, rRNA was depleted using the Pan-Prokaryote riboPOOLs kit (siTOOLs Biotech). In brief, 1 μg of total RNA was incubated for 10 min at 68 °C and 30 min at 37 °C with 100 pmol of rRNA-specific biotinylated DNA probes in 2.5 mM Tris-HCl pH 7.5, 0.25 mM EDTA, and 500 mM NaCl. DNA-rRNA hybrids were depleted from total RNA by two consecutive 15 min incubations with 0.45 mg streptavidin-coated magnetic Dynabeads MyOne C1 (ThermoFisher Scientific) in 2.5 mM Tris-HCl pH 7.5, 0.25 mM EDTA, and 1 M NaCl at 37 °C. The rRNA-depleted RNA samples were purified using the Zymo RNA Clean & Concentrator kit combined with DNase treatment on a solid support (Zymo Research). cDNA libraries were prepared using the NEBNext Multiplex Small RNA Library Prep kit for Illumina (NEB) in accordance with the manufacturers’ instructions.

Library preparation and sequencing took place at the Earlham Institute, and were processed by Newcastle University Bioinformatics Support Unit. Briefly, raw sequencing reads were checked using Fast QC, reads were mapped to *Bacteroides cellulosilyticus* DSM 14838 (GCA_000158035) downloaded from Ensembl (assembly ID ASM15803v1). Reads were quantified against genes contained in the Ensembl annotation using featureCounts from the Rsubread package^50^. Counts were normalized by Trimmed Median of Means (TMM) as implemented in DESeq2, and differentially expressed genes determined according to a Negative Binomial model as per DESeq2.

### Proteomic analysis of *D. gadei*

#### Sample preparation

*Dysgonomonas gadei* cells were suspended in 5% sodium dodecyl sulfate (SDS) in 50 mM triethylammonium bicarbonate (TEAB) pH 7.5. The samples were subsequently sonicated using an ultrasonic homogenizer (Hielscher) for 1 minute. The whole-cell lysate was centrifuged at 10,000 × *g* for 5 min to remove cellular debris. Protein concentration was determined using a bicinchoninic acid (BCA) protein assay (Thermo Scientific). A total of 20 μg protein was used for further processing. Proteins were reduced by incubation with 20 mM tris(2-carboxyethyl)phosphine for 15 min at 47 °C, and subsequently alkylated with 20 mM iodoacetamide for 30 minutes at room temperature in the dark. Proteomic sample preparation was performed using the suspension trapping (S-Trap) sample preparation method ^51^, as recommended by the supplier (ProtiFi™, Huntington NY). Briefly, 2.5 μl of 12% phosphoric acid was added to each sample, followed by the addition of 165 μl S-Trap binding buffer (90% methanol in 100 mM TEAB pH 7.1). The acidified samples were added, separately, to S-Trap micro-spin columns and centrifuged at 4,000 × *g* for 1 min until the solution has passed through the filter. Each S-Trap micro-spin column was washed with 150 μl S-trap binding buffer by centrifugation at 4,000 × *g* for 1 min. This process was repeated for a total of five washes. Twenty-five μl of 50 mM TEAB containing trypsin (1:10 ratio of trypsin:protein) was added to each sample, followed by proteolytic digestion for 2 h at 47 °C using a thermomixer (Eppendorf). Peptides were eluted with 50 mM TEAB pH 8.0 and centrifugation at 1,000 × *g* for 1 min. Elution steps were repeated using 0.2% formic acid and 0.2% formic acid in 50% acetonitrile, respectively. The three eluates from each sample were combined and dried using a speed-vac before storage at −80 °C.

#### Mass Spectrometry

Peptides were dissolved in 2% acetonitrile containing 0.1% trifluoroacetic acid, and each sample was independently analyzed on an Orbitrap Fusion Lumos Tribrid mass spectrometer (Thermo Fisher Scientific), connected to an UltiMate 3000 RSLCnano System (Thermo Fisher Scientific). Peptides were injected on a PepMap 100 C_18_ LC trap column (300 μm ID × 5 mm, 5 μm, 100 Å) followed by separation on an EASY-Spray nanoLC C_18_ column (75 μm ID × 50 cm, 2 μm, 100 Å) at a flow rate of 250 nl/min. Solvent A was water containing 0.1% formic acid, and solvent B was 80% acetonitrile containing 0.1% formic acid. The gradient used for analysis was as follows: solvent B was maintained at 2% for 5 min, followed by an increase from 2 to 35% B in 120 min, 35-90% B in 0.5 min, maintained at 90% B for 4 min, followed by a decrease to 3% in 0.5 min and equilibration at 2% for 10 min. The Orbitrap Fusion Lumos was operated in positive-ion data-dependent mode. The precursor ion scan (full scan) was performed in the Orbitrap in the range of 400-1,600 *m/z* with a resolution of 120,000 at 200 *m/z*, an automatic gain control (AGC) target of 4 × 10^5^ and an ion injection time of 50 ms. MS/MS spectra were acquired in the linear ion trap (IT) using Rapid scan mode after high-energy collisional dissociation (HCD) fragmentation. An HCD collision energy of 30% was used, the AGC target was set to 1 × 10^4^ and dynamic injection time mode was allowed. The number of MS/MS events between full scans was determined on-the-fly to maintain a 3 s fixed duty cycle. Dynamic exclusion of ions within a ±10 ppm *m/z* window was implemented using a 35 s exclusion duration. An electrospray voltage of 2.0 kV and capillary temperature of 275 °C, with no sheath and auxiliary gas flow, was used.

All mass spectra were analyzed using MaxQuant 1.6.12.0^52^, and searched against the *Dysgonomonas gadei* ATCC BAA-286 proteome database downloaded from Uniprot (accessed 09.01.2020). Peak list generation was performed within MaxQuant and searches performed using default parameters and the built-in Andromeda search engine^53^. The enzyme specificity was set to consider fully tryptic peptides, and two missed cleavages were allowed. Oxidation of methionine, N-terminal acetylation and deamidation of asparagine and glutamine were allowed as variable modifications. Carbamidomethylation of cysteine was allowed as a fixed modification. A protein and peptide false discovery rate (FDR) of less than 1% was employed in MaxQuant. Proteins that contained similar peptides and could not be differentiated based on MS/MS analysis alone were grouped to satisfy the principles of parsimony. Reverse hits, contaminants, and proteins only identified by site modifications were removed before downstream analysis.

### Generation of expression constructs

Unless stated otherwise genes were purchased as codon-optimized constructs from Twist Biosciences. *D. gadei* and *B. finegoldii* genes were cloned from genomic DNA using standard restriction cloning methods, followed by ligation into pET28a vectors and transformation into TOP10 cells (Novagen) with subsequent sequencing of selected purified recombinant plasmids by Eurofins for confirmation.

### Protein Expression and Purification

#### Expression and purification of Dg_GH4185a_, Dg_GH4185b_, Phage_GH4185_, Myxo_GH4185_, Dg_GH172a_, Dg_GH172b_, Dg_GH172c_, and Myc_GH172_

Recombinant proteins were expressed in competent *E. coli* Tuner cells (Novagen) using pET28a vectors generated as above. Cells were grown in LB media at 37 °C with shaking, until turbidity reached an OD_600_ of ~0.6, wherein expression was induced with 0.2 mM IPTG and cells were further cultured for 16 hours at 16 °C. Sonication was used to lyse cells in 20 mM Tris, pH 8.0, 200 mM NaCl.

Enzymes were purified using immobilized metal affinity chromatography on cobalt TALON resin. Proteins were dialyzed into 20 mM HEPES, pH 8.0, 150 mM NaCl buffer dialysis (Medicell). For crystallography proteins were purified further via size-exclusion chromatography (HiLoad 16/600 Superdex 200, GE Healthcare) in 20 mM HEPES, pH 8.0, 150 mM NaCl. Protein purity was ascertained by SDS-PAGE and protein concentrations were determined using Nanodrop spectroscopy (Thermofisher).

#### Purification of Rv3707c and MSMEG_2107

For protein Rv3707c and MSMEG_2107 expression, an aliquot of competent BL21 DE3 *Escherichia coli* was transformed with plasmid and plated on LB agar supplemented with 50 μg/mL kanamycin. One plate of bacteria was scraped to inoculate 1 L of modified Studier’s autoinduction media^54^. The bacteria were incubated at 37 °C with shaking (180 RPM) until OD_600_= ~0.6, whereupon the flasks were cooled on ice with agitation to 20 °C and then returned shake overnight at 20 °C. After induction, the cultures were pelleted at 3990 × *g* for 25 min at 4 °C. Pellets were resuspended in sterile PBS and pelleted at 7000 × *g* for 10 min. The supernatant was removed, and pellets were snap frozen in liquid nitrogen and stored at −20 °C until preparation.

Rv3707c was purified by suspension of one pellet in cold lysis buffer (25 mM HEPES pH8; 400 mM NaCl; 5% glycerol; 50 mM l-arginine; 50 mM l-glutamic acid; 1 mM beta-mercaptoethanol). 1 mg ml^−1^ deoxyribonuclease I from bovine pancreas (Sigma-Aldrich) was added to cell slurry and incubated for 30 minutes. Cells were lysed by three passages through a French pressure cell. Insoluble debris was then pelleted by centrifugation at 40,000 × *g* for 40 minutes (4 °C). The supernatant was then processed by immobilized metal affinity chromatography (IMAC) on a gravity column containing 2 mL bed volume of cOmplete His Tag purification resin (Roche). After loading the lysate, the column was washed with 80 mL of lysis buffer, then eluted with an imidazole gradient of 50, 100, 250, and 500 mM. Protein-containing fractions were pooled and dialyzed exhaustively against three liters of dialysis buffer (25 mM HEPES pH8; 400 mM NaCl; 5% glycerol; 50 mM l-arginine; 50 mM l-glutamic acid; 2 mM dithiothreitol). The crude protein was concentrated to a final volume of 0.5 mL on a 30k Da molecular weight cutoff Pierce protein concentrator (Thermo Scientific). This fraction was then further purified by size exclusion chromatography on an AKTA Prime system with a SuperDex 26/600 S200 column in the above dialysis buffer before being concentrated. The protein was always used freshly prepared.

#### Purification of Noc_GH172_ and Mab_GH4185_

One plate of BL21-DE3 transformed with an appropriate plasmid was used to inoculate 1L Terrific Broth supplemented with kanamycin. The culture was grown to an OD_600_ of 0.6 and induced with 0.25 mM IPTG and grown overnight at 20 °C. After harvest of biomass as in purification of Rv3707c, cell pellets were resuspended in a buffer containing 25 mM HEPES; 40 mM NaCl, lysozyme, and DNAse I. Subsequent purification steps were identical to those in Rv3707c, but in the above, simpler buffer, omitting lysozyme and DNAse I.

### Crystallography

#### Crystallography of Dg_GH172c_

Crystallization of Dg_GH172c_ sample at 10 mg ml^−1^ was screened using commercial kits (Molecular Dimensions and Hampton Research) with vapor diffusion sitting drop method. Crystals formed in a buffer containing 0.1 M MOPS/Sodium HEPES pH 7.5, 0.12 M alcohol and 30% EDO_P8K over a period of 2 weeks. Crystals were harvested and flash cooled in liquid nitrogen. X-ray diffraction data were collected at the synchrotron beamline I24t Diamond light source (Didcot, UK) at a temperature of 100 K. The data were integrated using XDS^55^ via XIA2 ^56^ and scaled with Aimless^57^. The space group was confirmed with Pointless. The phase problem was solved by molecular replacement using Phaser and PDB model 4KQ7 as search model from *Bacteroides uniformis*^58^. Subsequently, the structure was auto built with CCP4build on CCP4cloud^59^. The model was improved by rounds of manual building with COOT and refinement with Refmac^60,61^. The final model was validated with Molprobity^62^ and COOT^60^.

#### Crystallography of Rv3707c

Rv3707c was concentrated to 7 mg ml^−1^ and loaded into a sitting drop crystallization tray (Clover) using JCSG-plus crystallization screen (Molecular Dimensions) at a ratio of 2 μL protein 2 μL reservoir solution. Crystals were formed in a buffer containing 0.8 M succinic acid pH 7 over a period of 1 month. The crystals were cryoprotected in a solution of 0.8 M succinic acid pH 7 with 30% glycerol and flash cooled in liquid nitrogen. X-ray diffraction data were collected at beamline I04 of Diamond light source, Oxford. Data were auto-processed using Xia2 and general file manipulations were performed using the CCP4 suite of programmes^56,59^. Data were phased by molecular replacement using the Alphafold structural prediction of Rv3707c as a search model using PHASER (TFZ of 21.1)^58^. The structure was then auto built in PHENIX with rounds of refinement carried out by PHENIX-refine and manually in COOT^60,63^.

### Summary of methods for phylogenetic analyses

Two alignments were used to infer respectively a global phylogeny for members of the PF13810 family and a more restricted phylogeny focusing on close relatives to the functionally characterized proteins. The global alignment was derived from the “Full” Pfam alignment for PF13810 made of 1145 sequences and 1321 aligned sites (http://pfam.xfam.org/family/PF13810#tabview=tab3). This led to an alignment of 753 sequences and 179 aligned sites by: (i) deleting partial sequences that did not include the catalytic residues or that corresponded to obsolete sequences (nine sequences) and (ii) adding the three sequences from *Mycobacterium abscessus* (strain 4529 available at the Integrated Microbial Genomes & Microbiomes (IMG/M) database: 2635695100/Ga0069448_11324, 2635694794/Ga0069448_1118, 2635698039/Ga0069448_113269) and (iii) deleting sites made of a majority of indels.

For the restricted alignment 39 complete sequences were aligned, including the seven proteins investigated in this study - enzymatic characterization and one structure. The sequences were aligned with Clustal Omega using default settings in SEAVIEW v.4.6.4 ^64,65^. Following minor manual adjustments of the alignment the mask function of SEAVIEW was used to selected aligned residues that included conserved blocks of sequences with no more than two indels leading to 200 aligned sites. The DUF4185 alignment is available as Supplementary Dataset 1 for respectively the global and restricted alignment. IQ-TREE (v.1.6.12) was used to generate maximum likelihood phylogenies using automatic model selection^66^. The selected models were LG+F+I+G4 for the global alignment and WAG+I+G4 for the restricted alignment using the Bayesian Information Criterion (BIC). Bootstraps (100 replicates) were computed to assess branch reliability. iTOL (interactive tree of life) was used to generate the figures^67^. The global phylogeny was annotated with taxonomy information derived from the UniProt database (https://www.uniprot.org/)^68^.

### Protein structure prediction by AlphaFold 2.0

Amino acid sequences were submitted to the Colabfold_advanced.ipynb server for protein structure prediction^69^. Unless otherwise stated the default values were used, with the max_recycles set to 12. Figure coloring for pLDDT values were generated in ChimeraX 1.4 using the protein structure prediction tool and a custom palette.

### Timepoint assays

To assess enzymatic activity, reactions were initiated containing substrates (in water) and enzymes (in 20 mM HEPES pH 7.5, 150 mM NaCl, unless otherwise stated) of various concentrations, with 50 mM potassium phosphate buffer pH 7.2 as a dominant reaction buffer. Reactions were incubated at 37 °C for 16 hours and subsequently boiled to ensure enzymatic cessation. Time point samples were then analyzed using TLC or IC-PAD.

### Porous graphitic carbon chromatography clean-up of Rv3707c and MSMEG_2107 arabinogalactan hydrolysis assays

Due to the high concentration of L-arginine and L-glutamic acid in the buffer used for purification of Rv3707c and MSMEG_2107, enzyme assays were unsuitable for HPAEC-PAD analysis without prior solid phase extraction. To this end, at each timepoint, reaction mixtures were loaded onto a Hypersep Hypercarb SPE cartridge (Thermo Scientific) which had been washed with acetonitrile and 50% THF in water and exhaustively equilibrated with water prior to loading. Reaction products were then eluted in 80% acetonitrile in ddH2O with 0.1% trifluoracetic acid (Sigma-Aldrich) and dried by evaporation in a SpeedVac concentrator before being reconstituted in the original volume of water.

### Kinetic analysis of GH172 arabinofuranosidase activity

Enzymes (100 nM) were incubated with the indicated concentrations of pNP-α-d-Ara*f* or AG. Assays were performed in technical triplicate at 37 °C in 20 mM HEPES pH 7.5. for pNP, absorbances were measured at 400 nm and graphs were plotted in GraphPad Prism 9.3.1. For Ag, arabinose concentration was measured by IC-PAD (see below) with reference to a standard curve.

### Thin-layer chromatography (TLC)

Purified proteins were incubated with at 0.1-5 μM (as indicated) with substrates (for methanolysis, methanol was added to the reaction mixture at a final concentration of 10%) for 16 h at 37 °C to ensure reaction completion (unless otherwise indicated). Using TLC plate aluminum foils (Silicagel 60, 20 × 20, Merck) which were cut to the desired size (minimum height of 10 cm), these reaction samples were spotted (6 μl, unless otherwise indicated) and allowed to dry. TLC plates were run (twice) in solvent (1-butanol/acetic acid/ water 2:1:1 (v/v)). Plates were then removed and dried before visualization of sugars was obtained via immersion of TLC plate in Orcinol stain. Plates were dried and developed through heating between 50 °C and 80 °C.

### Ion Chromatography with Pulsed Amperometric Detection (IC-PAD)

Oligosaccharides from enzymatic polysaccharide digestion were analyzed using a CARBOPAC PA-300 anion exchange column (ThermoFisher) on an ICS-6000 system. Detection enabled by PAD using a gold working electrode and a PdH reference electrode with standard Carbo Quad waveform. Buffer A – 100 mM NaOH, Buffer B – 100 mM NaOH, 0.5 M Na Acetate. Each sample was run at a constant flow of 0.25 ml·min^−1^ for 100 minutes using the following program after injection: 0-10 min; isocratic 100% buffer A, 10-70 min; linear gradient to 60% buffer B, 70-80 min; 100% buffer B. The column was then washed with 10 mins of 500 mM NaOH, then 10 min re-equilibration in 100% buffer A. l-arabino-oligosaccharides (DP = 2-9) obtained commercially (Megazyme) were used as standards at a concentration of 25 μM. Data were processed using Chromeleon™ Chromatography Management System V.6.8. Final graphs were created using GraphPad Prism 8.0.1.

### Purification of mycobacterial arabinogalactan

Large scale purification of mycobacterial arabinogalactan was achieved by established methodologies^70^. In brief, 8 L of mycobacterial culture was grown to mid-exponential phase, cultures were pelleted and resuspended in a minimal volume of phosphate-buffered saline (VWR) and lysed using an Emulsiflex. The lysate was then boiled in a final concentration of 1% sodium dodecyl sulphate (SDS) and refluxed. Insoluble material (containing mycolyl-arabinogalactan-peptidoglycan complex) was collected by centrifugation and washed exhaustively with water to remove SDS. The mycolate layer was removed by saponification by KOH in methanol at 37 °C for 3 days. Cell wall material was then washed repeatedly to remove saponified mycolic acids with diethyl ether. The phosphodiester linkage between AG and PG was then cleaved by treatment with H2SO4 at 95 °C before being neutralized with sodium carbonate. The resultant solubilized arabinogalactan was collected in the supernatant, dialyzed exhaustively against water and lyophilized (yield= ~22.5 mg·L^−1^).

### Purification of d-arabinan

Arabinogalactan (5 mg/mL) was digested in a 10 mL total volume of 25 mM MOPS buffer pH 7. One μM final concentration of BACFIN_04787 and BACFIN_08810 were added, and the reaction was incubated at 37°C for 48h. An aliquot of the reaction was analyzed by TLC to verify the hydrolysis of the substrate and galactose release. Then, the sample was dialyzed over-night against 5 L of deionized water using a 1 kDa membrane, to eliminate residual galactose from the reaction mixture. The sample was then freeze-dried and resuspended in 0.4 mL water. A TLC analysis confirmed the elimination of the residual galactose from the sample, and this was further confirmed through acid hydrolysis of the resulting D-arabinan to determine total abundance. To quantify the purity of isolated d-arabinan, an aliquot of both (0.25 mg/mL) d-arabinan and arabinogalactan were treated by acid hydrolysis using 300 mM HCl at 100°C for 1h. Samples were neutralised to pH 7 with NaOH and analysed by HPAEC. Quantitation was based on the migration of standards and the ratio between galactose and arabinose.

### Purification of mycobacterial lipoglycans

One liter of mycobacterial culture was grown to mid exponential phase and pelleted as previously. The pellet was resuspended in 20 ml PBS, 0.1% Tween-80, chilled and lysed by bead-beating. Lysate was transferred to a Teflon-capped glass tube and vortexed with an equal volume of citrate buffer saturated with phenol (Sigma-Aldrich), and heated to 80 °C for 3 h, vortexing every hour. A biphase was generated by centrifugation at 2000 RPM at 10 °C, and the upper aqueous phase transferred to a fresh glass tube and hot phenol wash repeated twice more. The resultant protein free glycan mixture was dialyzed exhaustively against tap water overnight to remove trace phenol and lyophilized, yielding 34 mg of crude lipoglycans (LAM, LM, PIMS) per liter of culture.

### *Pseudomonas aeruginosa* pilin oligosaccharide extraction

Pilins were purified as described by Burrows and colleagues, with some modifications^42^. Briefly, *Pseudomonas aeruginosa* PA7 were streaked out in a grid pattern onto LB agar plates and grown for 24 hours at 37 °C. Cells were then scraped from all plates using sterile cell scrapers and resuspended in 4 ml of sterile phosphate-buffered saline (pH 7.4) per plate.

Pili were then sheared from the cell wall by vigorous vortexing of resuspended cells for 2 min. This suspension was centrifuged for 5 min at 6000 × *g*. The supernatant was centrifuged for 20 min at 20,000 × *g*. Supernatants were transferred to fresh tubes and MgCl_2_ was added to give a final concentration of 100 mM. Following inversion of these tubes to ensure mixing, samples were incubated at 4 °C overnight, allowing precipitation of sheared proteins. Samples were then centrifuged for 20 minutes at 20,000 × *g*, yielding a precipitate smudge which was resuspended in 50 mM NH_4_HCO_3_, pH 8.5 and transferred to fresh Eppendorf tubes. This solution was then dialyzed into the same buffer to eliminate excess MgCl_2_.

Bradford assays were then performed to assay the mass of protein in the sample, followed by digestion of the intact protein pilins using proteinase K in a 50:1 pilin to enzyme ratio by mass for 24 hours in the presence of 2 mM CaCl_2_. Glycans were then purified from digested proteins using porous graphitized carbon chromatography, where sugars were eluted from the column using a twofold increasing concentration series of a butan-1-ol:H_2_O gradient from 1:32 to 1:1 using 1 mL elutions^71^. Thin-layer chromatography (TLC) of eluates showed various oligomers of arabinan present in all fractions, all of which were subsequently used as substrates for potential arabinanases.

### Synthesis of pNP-α-d-arabinofuranoside

Para-nitrophenol (pNP)-α/β-d-arabinofuranoside synthesis was achieved following the established procedures ^23^.

### SEC-LS

Molecular weights were determined by size exclusion chromatography coupled light scattering using an Agilent MDS system with either an Agilent BioSEC 5 1000 Å, 4.6 × 300 mm, 5 μm or GE Superdex 200 5 15 mm columns at appropriate flow rates. Detector offsets were calibrated using a BSA standard and concentrations were determined by refractive index.

### Protein sequence alignment and structure visualization

Sequence alignments were performed using Clustal Omega and visualized using ESPript^64,72^. Figures of protein structures were generated with ChimeraX 1.4^73^.

### Fluorescent-conjugated mAGP hydrolysis assay

Fluorescently labeled mycolyl-arabinogalactan-peptidoglycan complex (mAGP) was isolated from *Corynebacterium glutamicum ATCC13032* following previously reported methods^36–38^. In brief, cells were grown in the presence of 5-AzFPA to saturation, reacted with DBCO-conjugated AF647 and then the mAGP was isolated. The isolated product was suspended in 2% SDS in PBS and split into the outlined treatment groups in Eppendorf tubes. Samples were pelleted by centrifugation at 15,000 × *g* for 5 min at 4°C then washed with PBS once (100 μL). Following this wash, the mAGP was resuspended in 90 μL PBS and enzyme stock added for a final concentration of 5 μM of each enzyme. Samples were incubated at 37 °C with rotation for 16 h. Following incubation, samples were pelleted by centrifugation at 15,000 × *g* for 5 min at 4 °C then washed with PBS twice (100 μL). The pellets were then suspended in 2% SDS in PBS, transferred to a 96-well plate and the fluorescence emission of each well was then measured on a Tecan Infinite M1000 Pro microplate reader. Monitoring of AF647 fluorescence was achieved by exciting at 648 nm ± 5 nm and detecting at 671 nm ± 5 nm. Z-position was set to 2 mm, and the fluorimeter gain was optimized and then kept constant. Data are reported in relative fluorescence units (RFU) normalized to PBS treated controls.

## Abbreviations

AG: arabinogalactan
DUF: domain of unknown function
GH: glycoside hydrolase
IC-PAD: ion chromatography/pulsed amperometric detection
LAM: Lipoarabinomannan
PUL: polysaccharide utilization locus

## Acknowledgements

We thank members of the Birmingham mycobacteriology group and the Newcastle glycobiology group for support and discussions. We gratefully acknowledge Diamond synchrotron source for access to X-ray beamtime.

## Funding

PJM is supported by a BBSRC David Phillips Fellowship (BB/S010122/1) and a BBSRC Impact Acceleration Award (1544084). ECL is supported by an Academy of Medical Sciences Springboard Award (SBF006\1048) and a Royal Society Research Grant (RGS\R2\202228). ALL is supported by a Wellcome Trust Investigator Award in Science (209437/Z/17/Z). SJW is supported by the Australian Research Council (DP210100233, DP210100233). STB is supported by a studentship from the Wellcome Trust. NPB is supported by a BBSRC studentship (BB/M011186/1). OA-J is funded by a BBSRC studentship (BB/M011186/1). JM-M was supported by an Advanced Grant from the European Research Council (Grant no. 322820) awarded to Harry J Gilbert. AC is funded by Academy of Medical Sciences Springboard (SBF005\1065) and a Royal Society research grant (RGS\R2\212050). This research was supported by the National Institute of Allergy and Infectious Disease (R01 Al-126592) and the NIH Common Fund (U01GM125288).

## Author Contributions

**Omar Al-Jourani** – Methodology, validation, formal analysis, investigation, data curation, visualization. **Samuel Benedict** – Methodology, validation, formal analysis, investigation, data curation, visualization, resources. **Jennifer Ross** – Methodology, validation, formal analysis, investigation, data curation, writing – original draft preparation, writing – review and editing. **Abigail Layton** – Methodology, validation, resources. **Phillip van der Peet** – Methodology, validation, formal analysis, resources. **Victoria Marando** – Methodology, formal analysis, investigation, visualization, writing – review and editing. **Nicholas P Bailey** – Methodology, validation, formal analysis, visualization. **Tiaan Heunis** – Methodology, formal analysis, resources. **Joseph Manion** – Investigation. **Francesca Mensitieri** – Investigation. **Aaron Franklin** – Investigation, resources. **Javier Abellon-Ruiz** – Investigation. **Sophia L Oram,** – Investigation. **Lauren Parsons** – Investigation. **Alan Cartmell** – Resources, supervision. **Gareth SA Wright** – Resources, supervision. **Arnaud Baslé** – Methodology, validation, writing – review and editing. **Matthias Trost** – Resources, supervision. **Bernard Henrissat** – Resources. **Jose Munoz-Munoz** – Investigation.**Robert P Hirt** – Investigation, resources, writing – review and editing, supervision. **Laura L. Kiessling** – Resources, supervision, funding acquisition, writing – review and editing. **Andrew Lovering** – Methodology, validation, formal analysis, writing – review and editing, supervision, funding acquisition. **Spencer J. Williams** – Resources, validation, formal analysis, writing – review and editing, supervision, funding acquisition. **Elisabeth C. Lowe** – Conceptualization, methodology, validation, formal analysis, data curation, writing – original draft preparation, writing – review and editing, visualization, supervision, funding acquisition. **Patrick J. Moynihan** – Conceptualization, methodology, validation, formal analysis, data curation, writing – original draft preparation, writing – review and editing, visualization, supervision, funding acquisition.

## Competing Interests

Drs. Moynihan and Lowe are co-inventors on an unpublished patent application pertaining to some of the enzymes described in this manuscript.

## Notes

### Competing Interest Statement

The authors have declared no competing interest.

### Summary of Updates

Text updated for readability.

