## Supplemental Figures and Tables for "Mining the human gut microbiome identifies mycobacterial d-arabinan degrading enzymes"

### Supplementary Figures

Figure S1: Identification of PULs associated with D-galactan degradation
Figure S2: *D. gadei* whole cell proteomics
Figure S3: TLC of GH172 enzyme catalysed reactions.
Figure S4: Michaelis-Menten Kinetics of GH172 enzymes against pNP-α-D-ara*f* and AG.
Figure S5: Hexameric assembly of GH172 enzymes
Figure S6: SEC-MALs of GH172 enzymes
Figure S7: Sequence alignment of GH172 enzymes
Figure S8: SEC-MALS analysis of $Dg_{GH172c}$ mutants
Figure S9: Phylogeny of DUF4185 enzymes found in selected acid-fast bacteria
Figure S10: Organisation of phage lysis cassettes in cluster C, DQ and a *Rhodococcus* phage singleton
Figure S11: TLC of DUF4185 enzyme catalysed reactions.
Figure S12: Sequence alignment of DUF4185 proteins
Figure S13: Start site analysis for Rv3707c
Figure S14: Rv1754c AlphaFold Prediction
Figure S15: Release of fluorescently labelled D-arabinan by selected acid-fast enzymes
Figure S16: Multiple catalytic residue conformations are predicted for Rv3707c

### Supplementary Tables

Table S1: Whole cell proteomics of *D. gadei* grown on D-arabinan, ordered by iBAQ. Proteins from PUL42 are coloured blue.
Table S2: Data collection and refinement statistics for the crystal structures of $Dg_{GH172c}$ and RV3707c
Table S3: Recipe for 2x defined minimal media

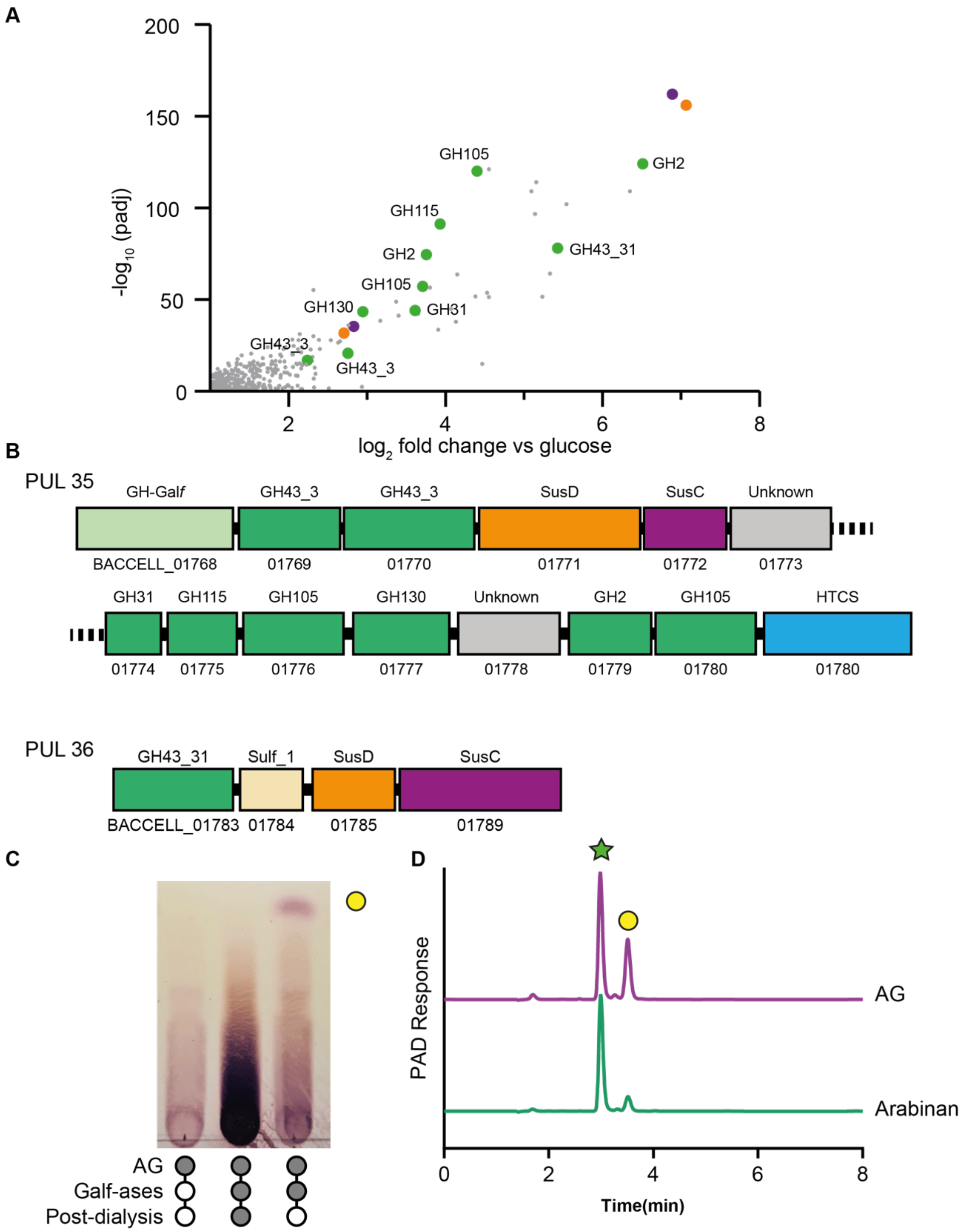


**Figure S1. Identification of PULs associated with D-galactan degradation. A)** Proteomic analysis of *B. cellulosilyticus* cells grown on AG as compared to those grown on glucose as a sole carbon source. **B)** PULs identified as being upregulated during growth on AG based on the proteomic analysis. **C)** AG from *M. smegmatis* was treated with 1 μM *B. finegoldii* enzymes BACFIN_08810 and BACFIN_04787 overnight, then dialysed to remove free galactose. **D)** Acid hydrolysed aliquots of treated arabinan and untreated arabinogalactan were analysed by HPAEC-PAD. Comparison to arabinose and galactose standards showed an approximate 70% reduction in galactose in the treated D-arabinan. Green star – D-arabinose, yellow circle – D-galactose.

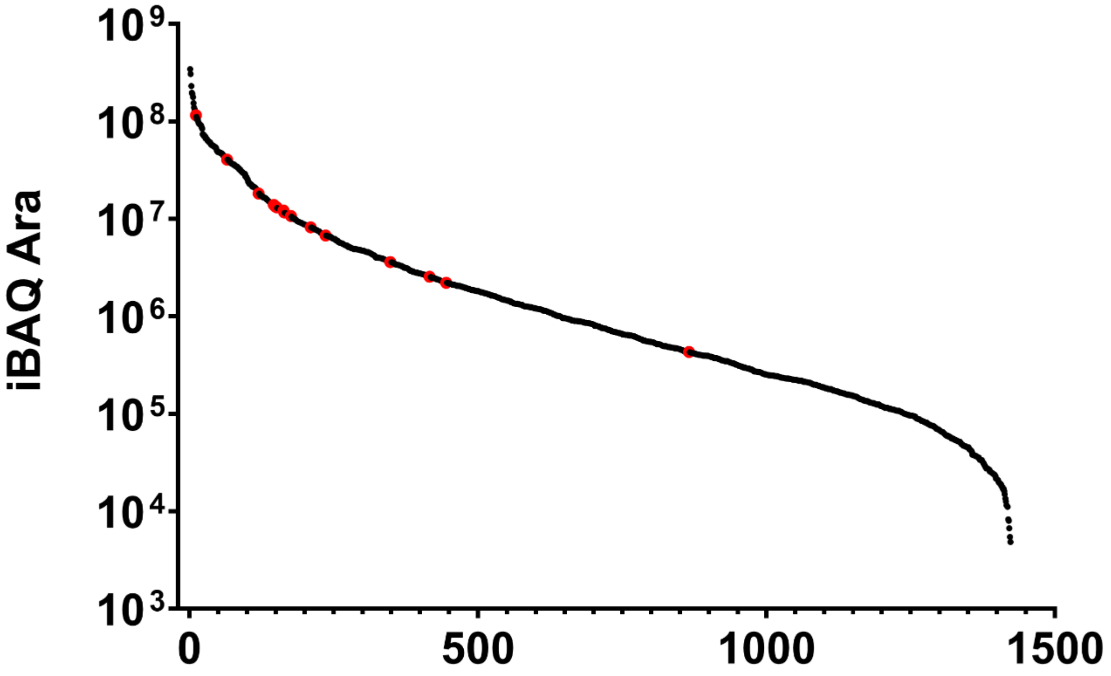


**Figure S2: *D. gadei* whole cell proteomics** Dynamic range of the *G. gadei* proteome, based on ranked iBAQ intensity, when grown on D-arabinan. Proteins from PUL42 are identified in red.

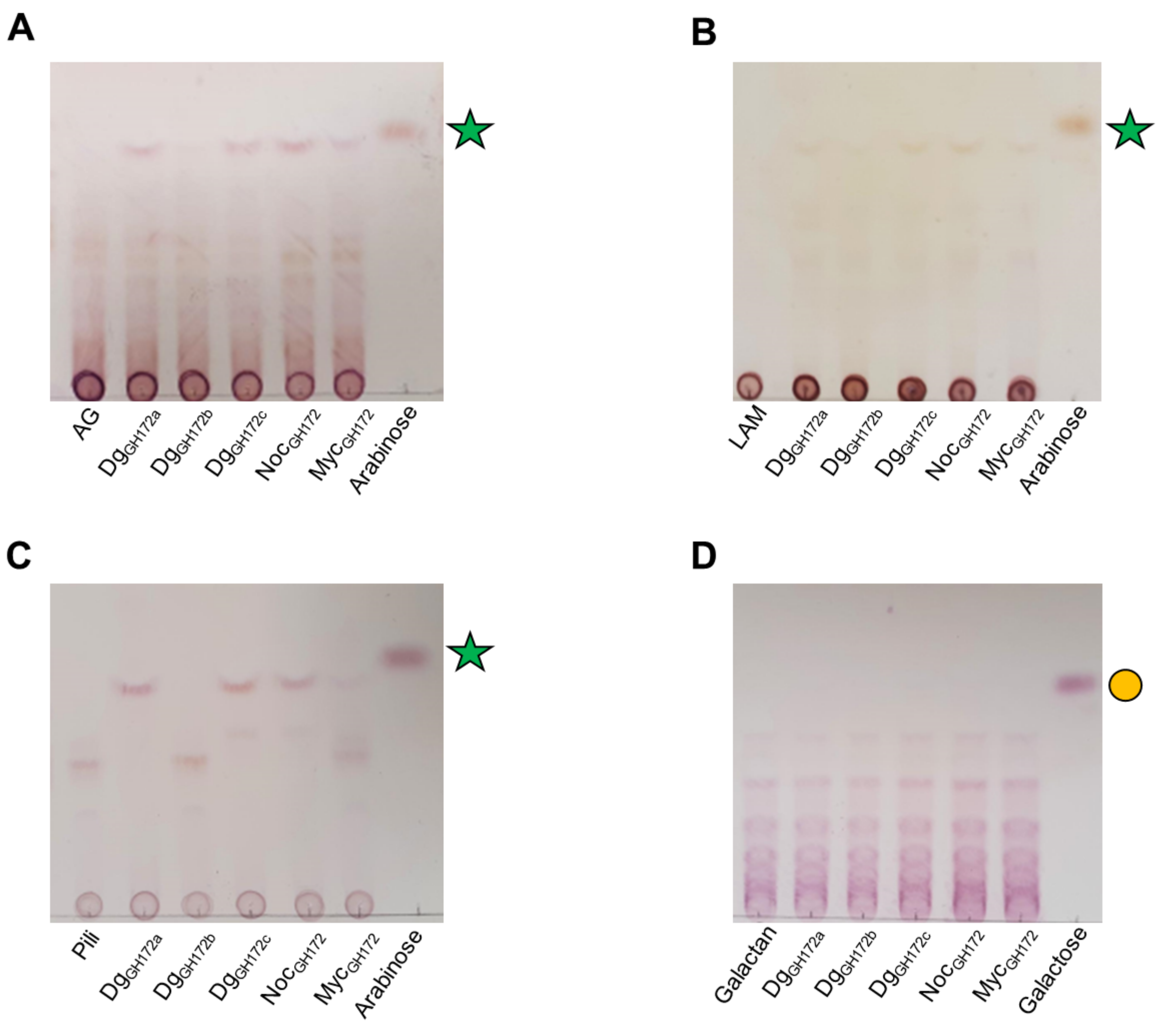


**Figure S3: TLC of GH172 enzyme catalysed reactions.** GH172 enzymes ($Dg_{GH172a}$, $Dg_{GH172b}$, $Dg_{GH172c}$, $Noc_{GH172}$, and $Myc_{GH172}$) were incubated with arabinogalactan (**A**), LAM (**B**), pilin oligosaccharides (**C**), and galactan (**D**) overnight at 37 °C, analysed by TLC and stained with orcinol.

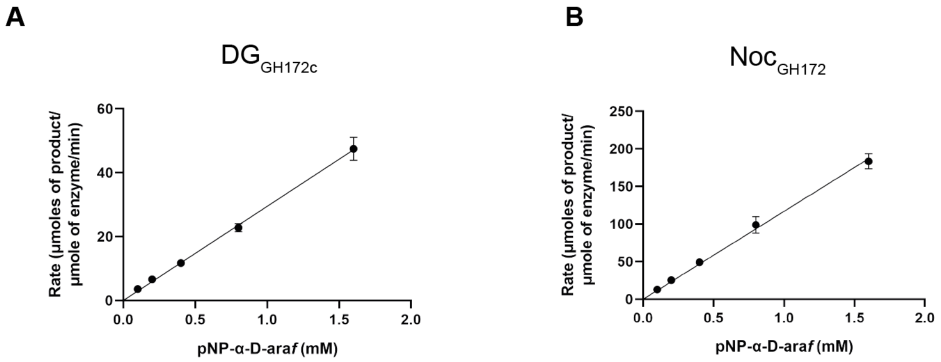


**Figure S4: Michaelis-Menten Kinetics of GH172 enzymes against pNP-α-D-Ara*f*.** Rates were measured with 100 nM enzyme at 37 °C in 20 mM HEPES pH 7.5, against varying concentrations of p-nitrophenyl-α-D-Ara*f* at 400 nm. Each concentration was repeated in technical triplicate, in some cases the error bars fall within the plotted data-points and are not visible in the plots.

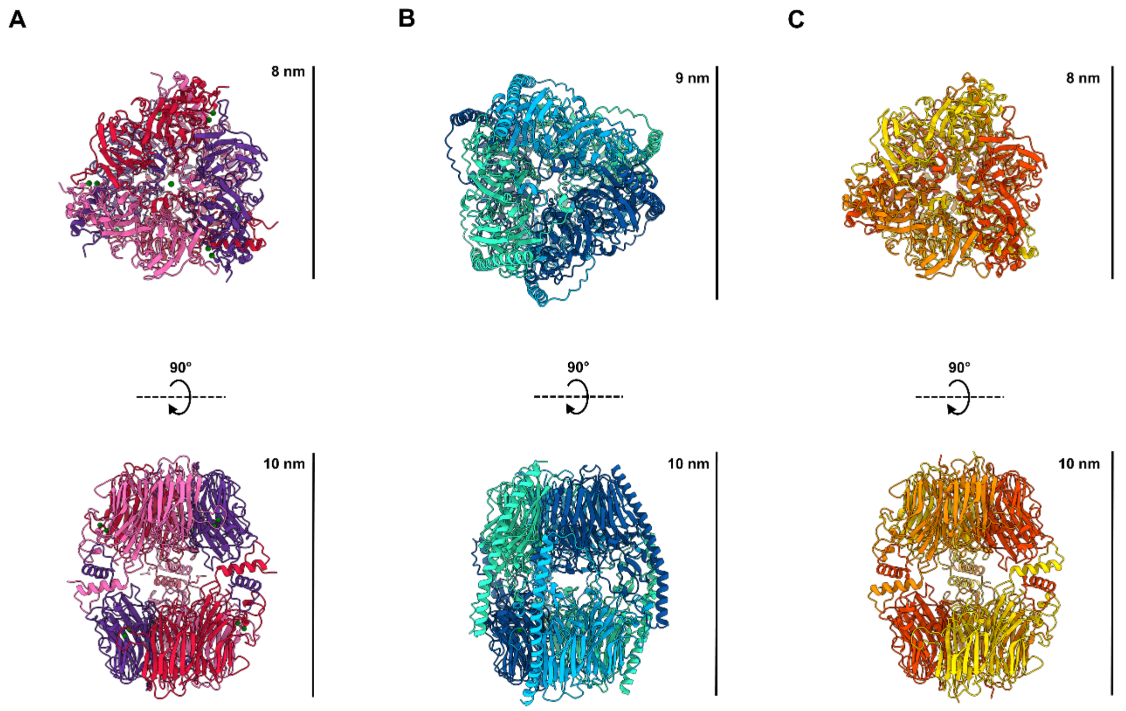


**Figure S5: Hexameric assembly of GH172 enzymes.** The biological, hexameric assembly of $Dg_{GH172c}$ (**A**) shown with secondary structure cartoon representation. The published structures of *B. dentium* (**B**, $Bd_{GH172}$, PDB: 7V1V) and *B. uniformis* (**C**, $Bu_{GH172}$, PDB: 4KQ7) are shown for comparison. Bound calcium ions are shown as green spheres.

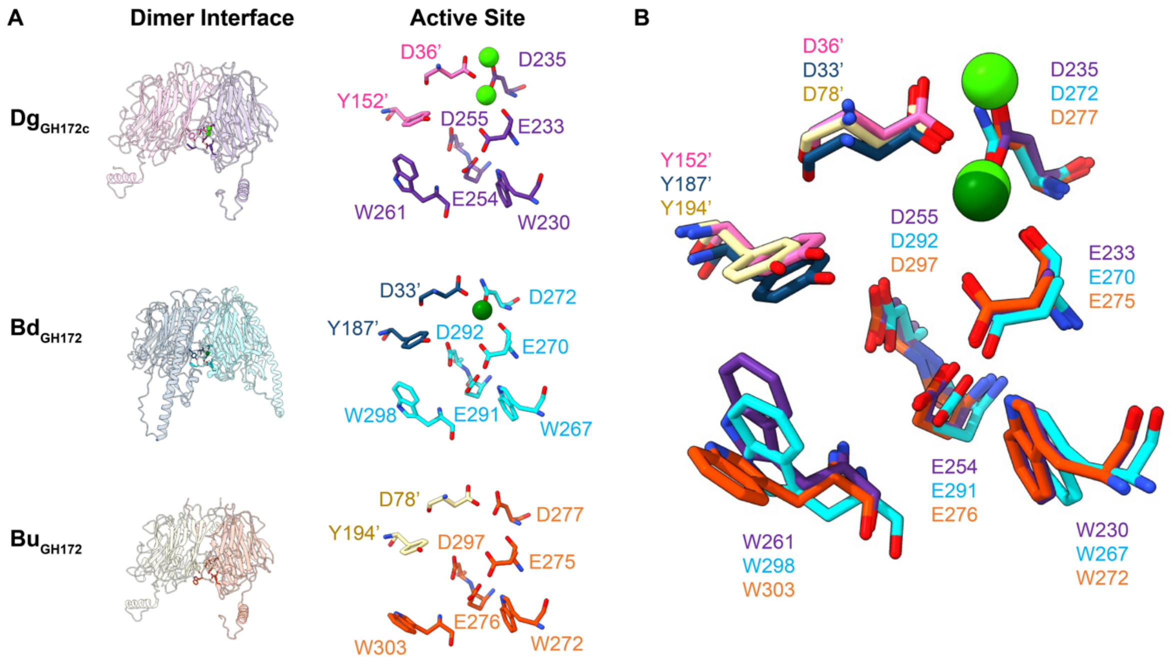


**Figure S6: Active site architecture of GH172 enzymes.** The GH172 active site is found at the interface of each pair of dimers (**A**) as shown for *D. gadei* ($\text{Dg}_{\text{GH172c}}$, PDB: 8AH3)*, B. dentium* ($\text{Bd}_{\text{GH172}}$, PDB: 7V1V) and *B. uniformis* ($\text{Bu}_{\text{GH172}}$, PDB: 4KQ7). Residues contributed from separate protomers are shown in a different colour. The active-site residues occupy equivalent positions in their respective structures (**B**; colouring as in A). Bound calcium ions are shown as green spheres.

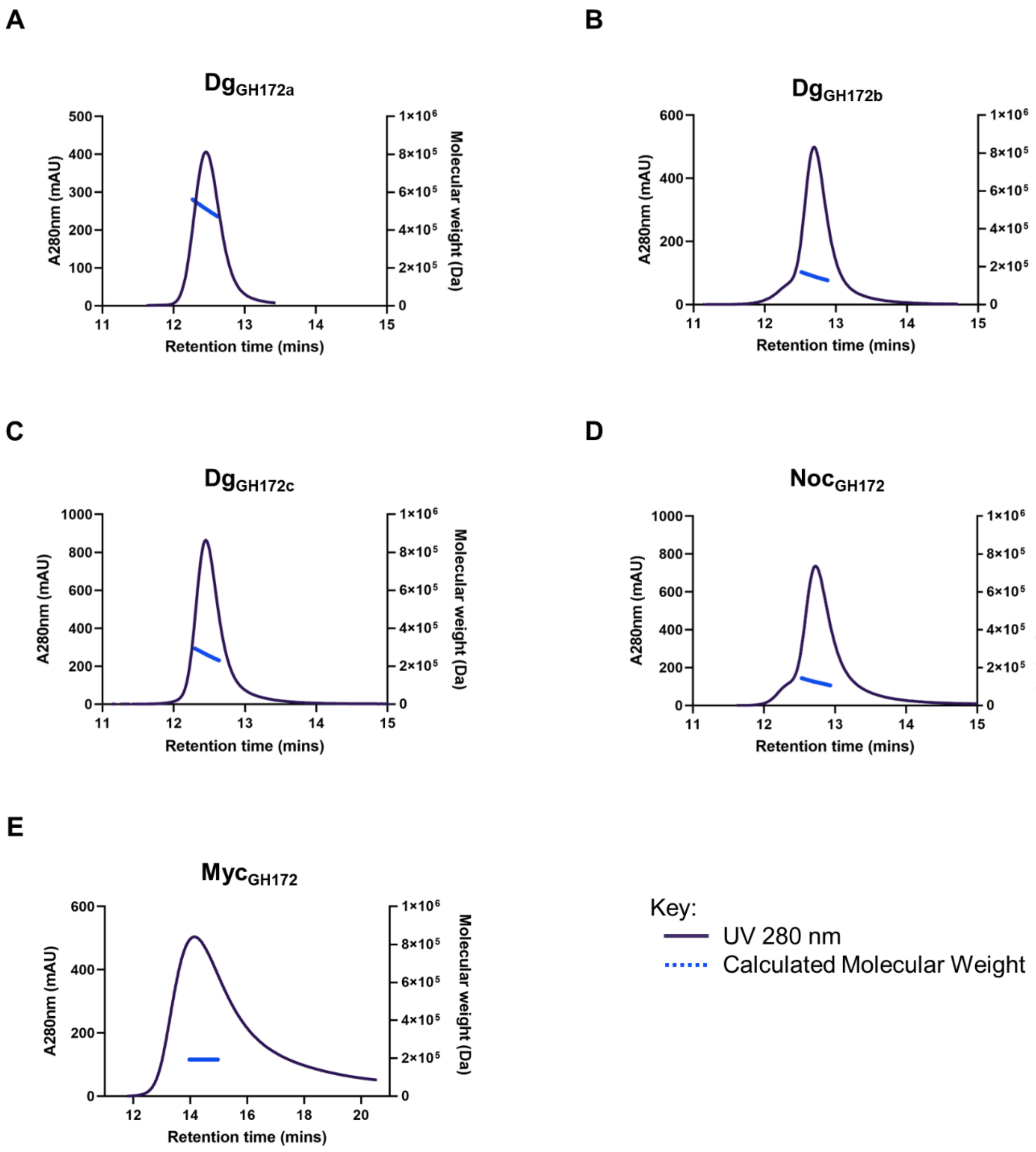


**Figure S7: SEC-MALs of GH172 enzymes.** SEC-MALS analysis of $Dg_{GH172a}$ (**A**), $Dg_{GH172b}$ (**B**), $Dg_{GH172c}$ (**C**), $Noc_{GH172}$ (**D**), and $Myc_{GH172}$ (**E**) allowing assignment of average masses to each GH172 enzyme (**Table 3**). Masses are consistent with: $Dg_{GH172a}$ being a dodecamer; $Dg_{GH172b}$ as a dimer species; $Dg_{GH172c}$ as a hexamer (consistent with its crystal structure); and $Noc_{GH172}$ and $Myc_{GH172}$ as trimeric species.

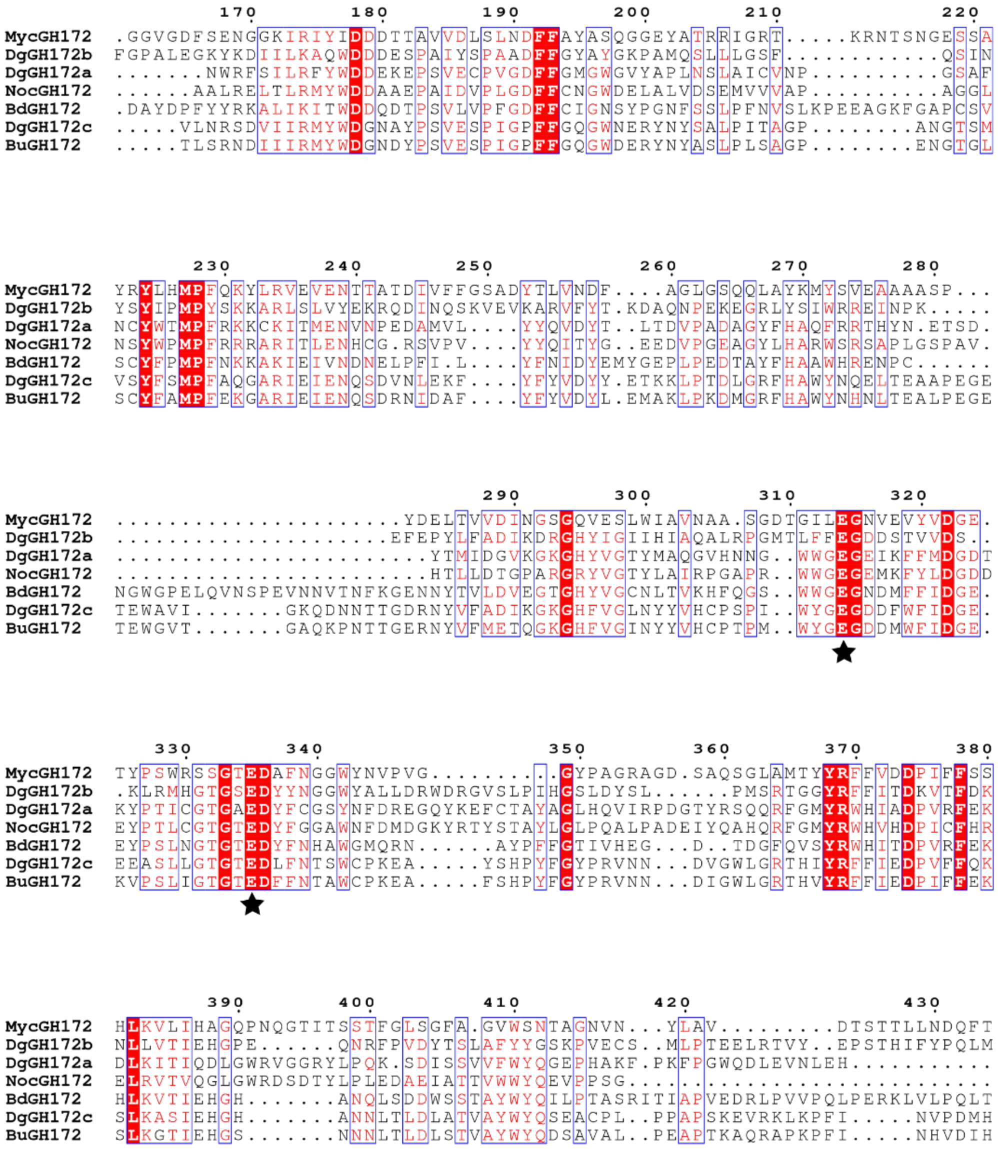


**Figure S8: Sequence alignment of GH172 enzymes.** Protein sequence alignment of GH172 enzymes ($Dg_{GH172a}$, $Dg_{GH172b}$, $Dg_{GH172c}$, $Bu_{GH172}$, $Bd_{GH172}$, $Noc_{GH172}$, and $Myc_{GH172}$). Catalytic residues have been highlighted by a black star. Red boxes with a white character show strict identity, and red character with blue frames show similar residues. Protein sequences were sourced from Uniprot and alignment was performed using Clustal Omega, and then formatted with ESPript 3.

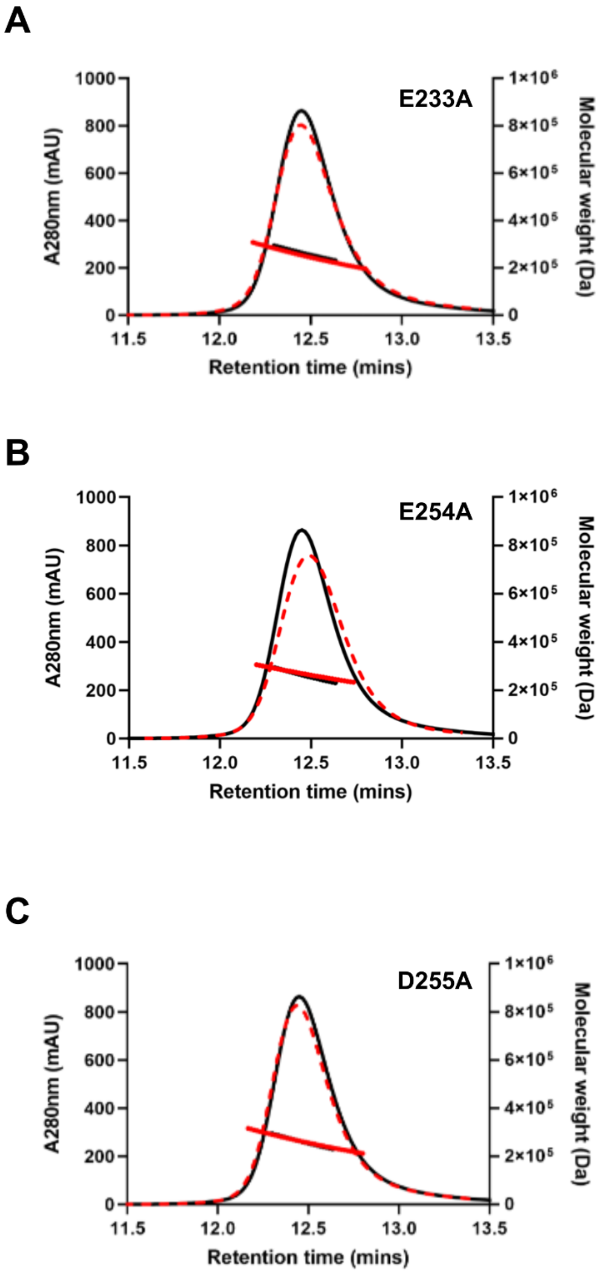


**Figure S9: SEC-LS analysis of Dg$_{GH172c}$ mutants.** SEC-MALS analysis of Dg$_{GH172c}$-E233A (**A**, dashed red line), Dg$_{GH172c}$ -E254A (**B**, dashed red line) and Dg$_{GH172c}$ -D255A (**C**, dashed red line) overlaid with Dg$_{GH172c}$-WT (black line, **A**/**B**/**C**). All mutants have a weight consistent with that of a hexamer and similar to WT.

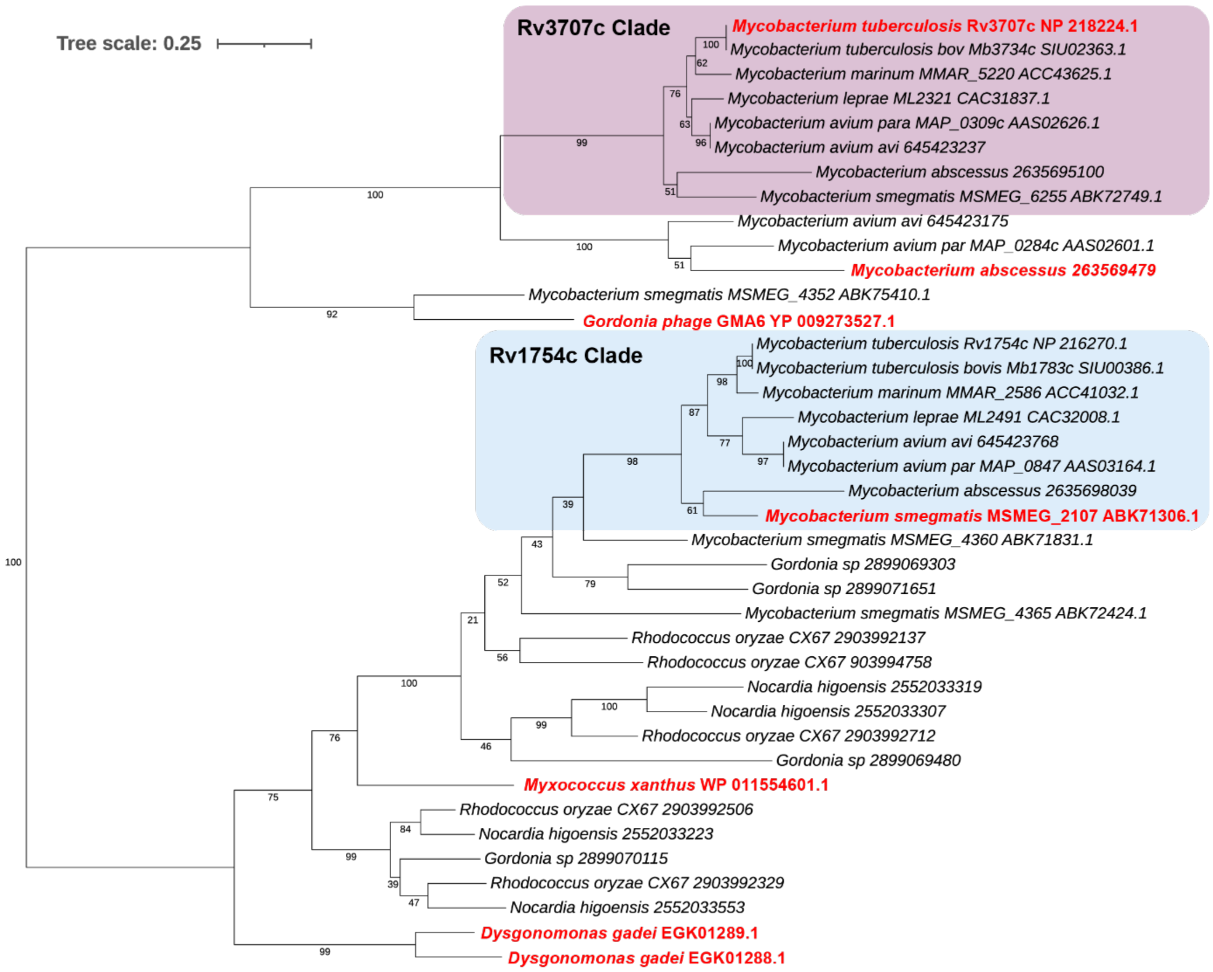


**Figure S10: Phylogeny of DUF4185 enzymes found in selected acid-fast bacteria**. A restricted alignment of 39 complete sequences from a selection of acid-fast bacteria and enzymes characterised in this study were aligned with Clustal Omega using default settings in SEAVIEW v.4.6.4 with some manual adjustments[58,59]. The alignment shows two clear clades of DUF4185 enzymes in acid fast bacteria with some further diversification, mainly in non-mycobacterial species.

.

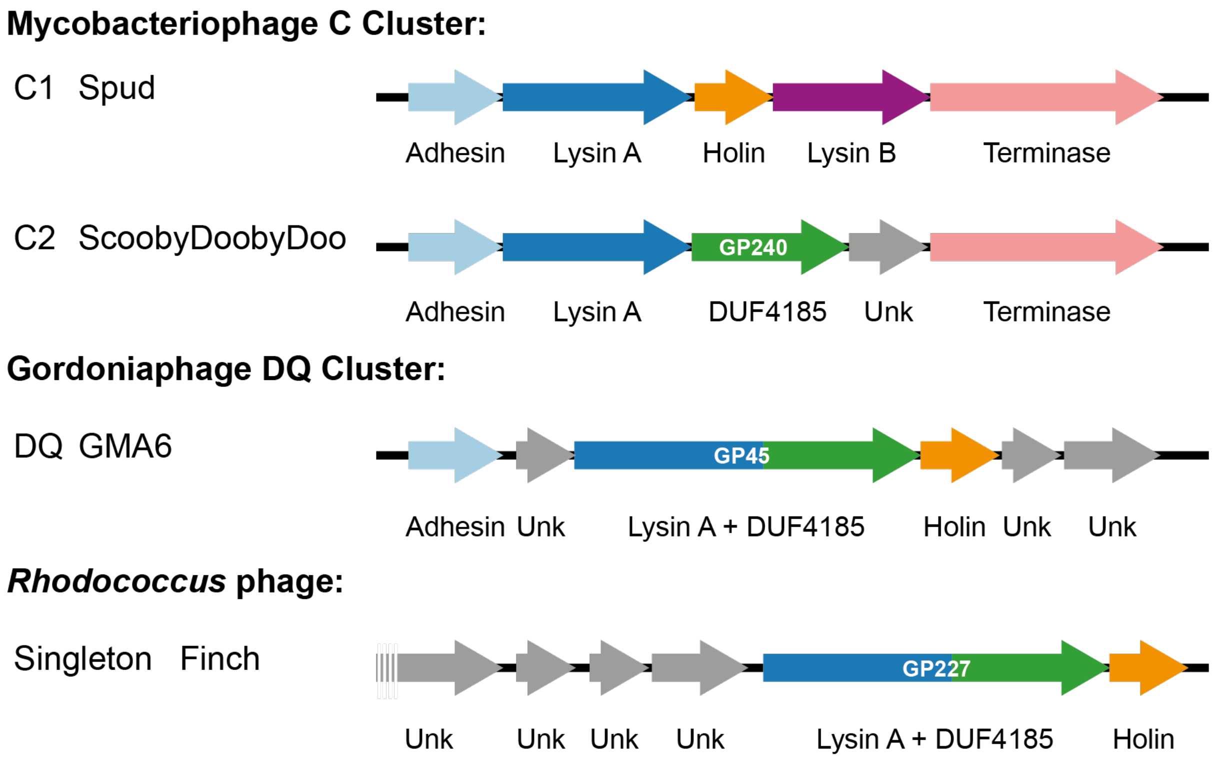


**Figure S11: Organisation of phage lysis cassettes in cluster C, DQ, and a *Rhodococcus* phage singleton**. Genomic regions encoding the lysis cassettes for clusters C1, C2, DQ in addition to the singleton Finch. Lysin B is absent in C, DQ and Finch and appears to be replaced by either a separate DUF4185 enzyme or as a LysinA-DUF4185 fusion.

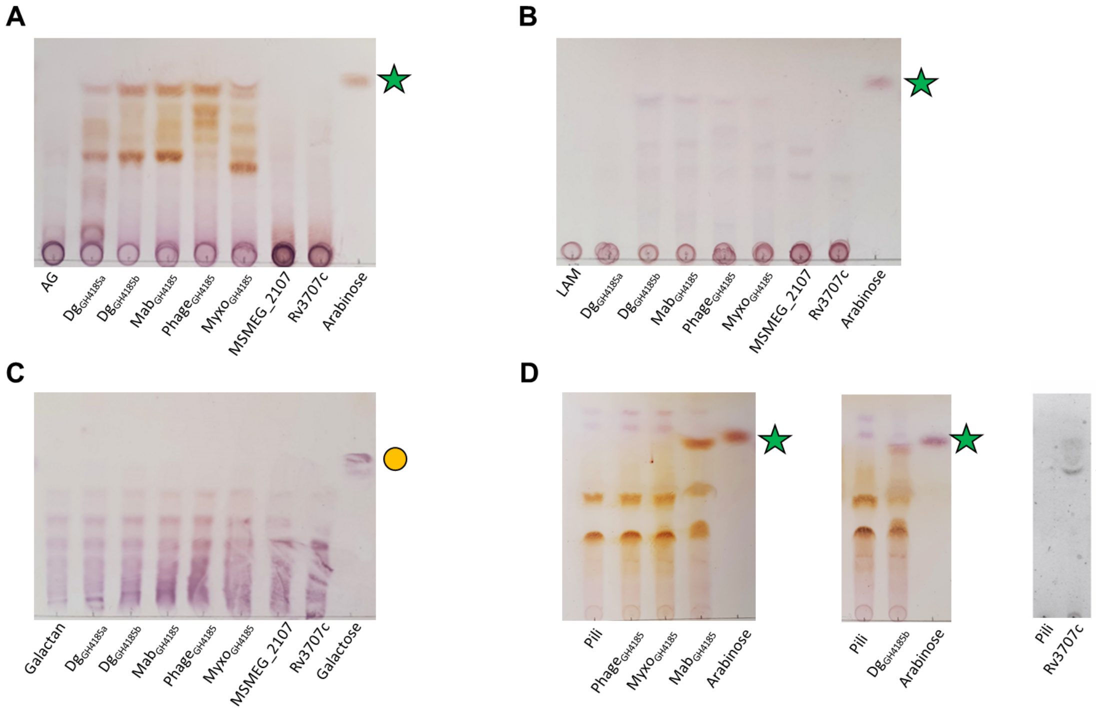


**Figure S12: Thin-layer chromatography (TLC) analysis of DUF4185 enzyme catalysed reactions.** DUF4185 enzymes ($Dg_{GH4185a}$, $Dg_{GH4185b}$, $Mab_{GH4185}$, $Phage_{GH4185}$, $Myxo_{GH4185}$, MSMEG_2107, and Rv3707c) were incubated with arabinogalactan (**A**), LAM (**B**), galactan (**C**), and pilin oligosaccharides (**D**) overnight at 37°C. 10 µL of each reaction was spotted at the origin of the TLC and the TLC was developed in a system consisting of 2:1:1 (v/v) of 1-butanol:acetic acid:water and visualised by staining with orcinol and charring. Yellow circles = D-galactose, green stars – D-arabinose.

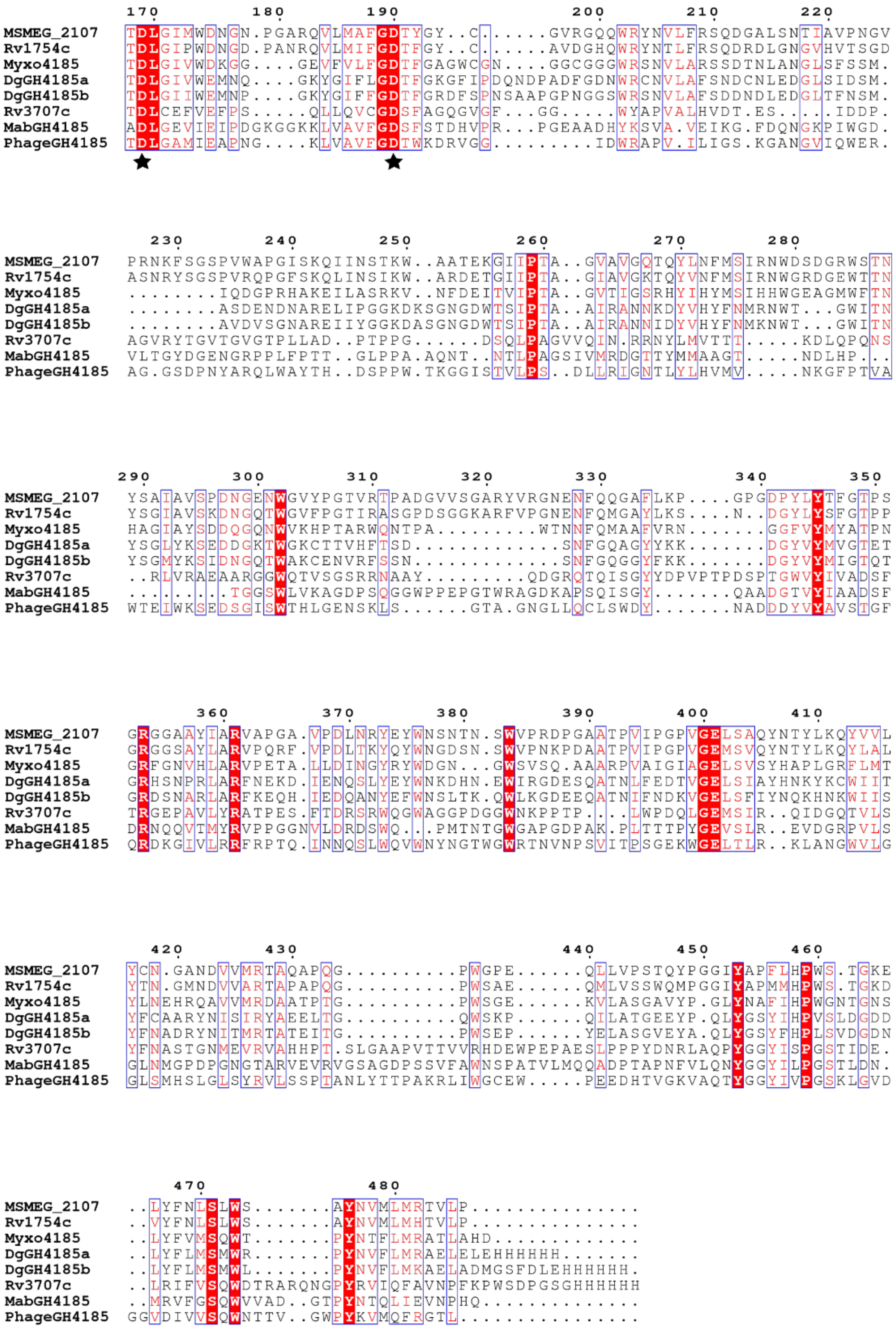


**Figure S13: Sequence alignment of DUF4185 proteins**

Protein sequence alignment of DUF4185 enzymes with putative active site residues highlighted with black stars. Red boxes with a white character show strict identity, and red character with blue frames. Sequence aligned using Clustal Omega and visualised using ESPript.

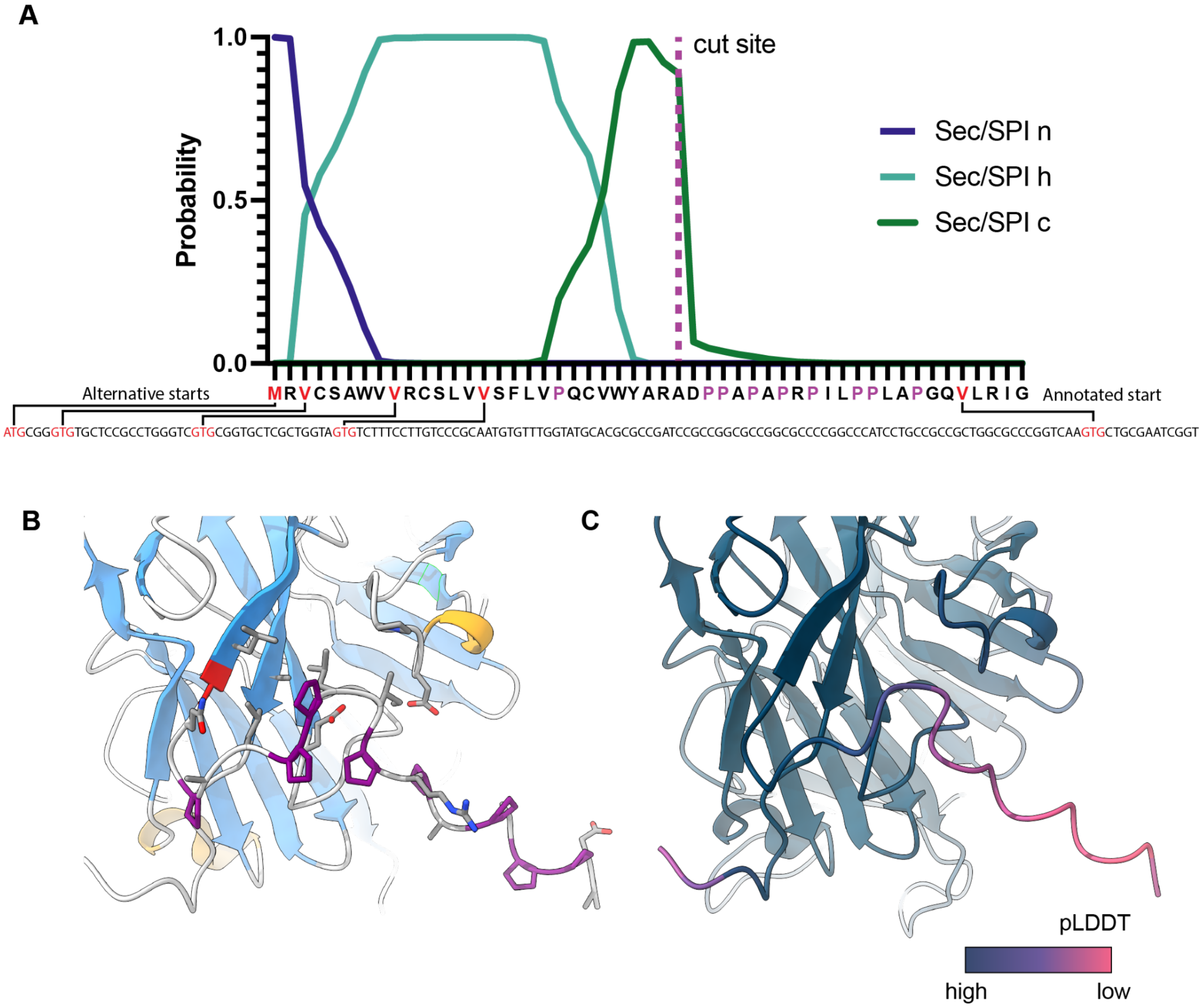


**Figure S14: Start site analysis for Rv3707c. A)** SignalP 6.0 output for Rv3707c with four potential upstream start sites included. Proline residues are indicated in purple. **B**) Close-up of N-terminus of AlphaFold 2.0 output for Rv3707c from the predicted cleavage point onward. The annotated start site is indicated in red and added prolines in purple, part of the unannotated N-terminus is predicted to form an ordered motif with high confidence by Alphafold 2.0 (**C**).

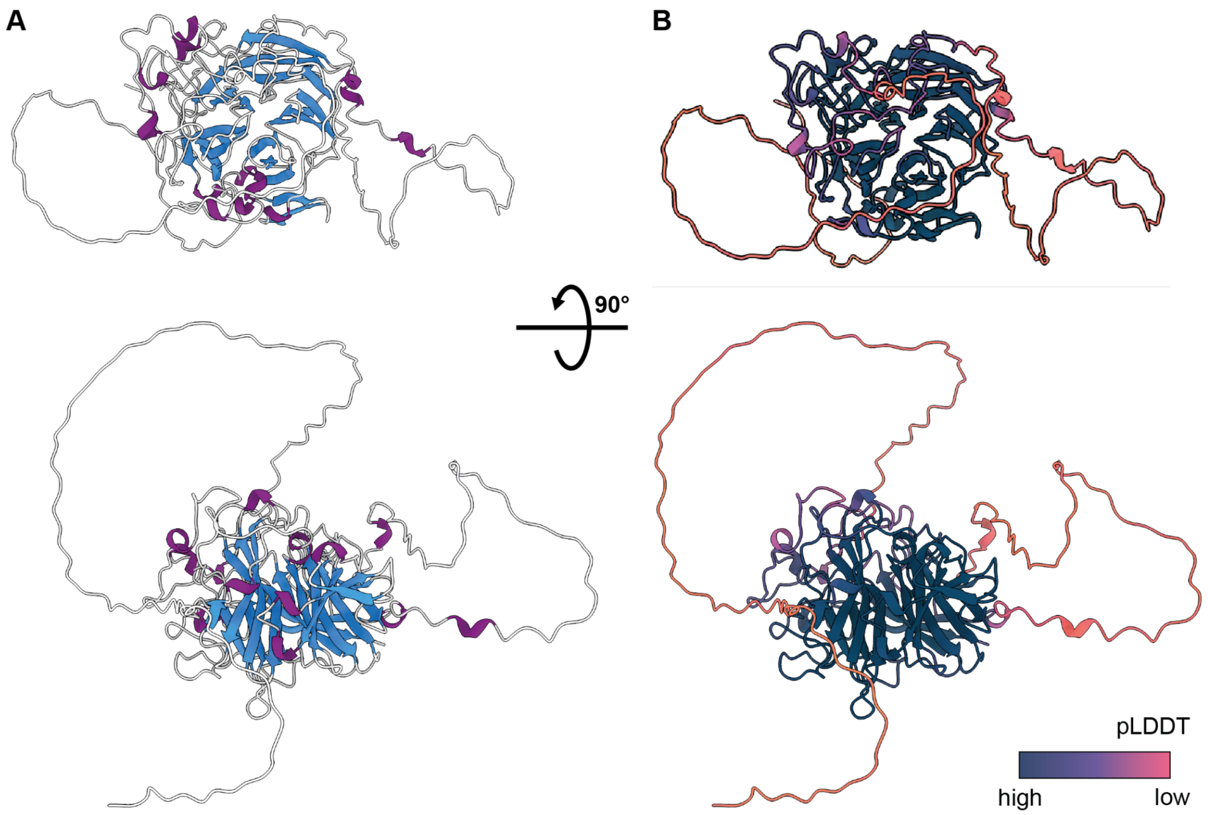


**Figure S15: Rv1754c AlphaFold Prediction.** Cartoon representation of Rv1754c as predicted by Alphafold coloured by secondary structure **(A)** and pLDDT **(B)**. AlphaFold prediction was computed with the default settings and 24 recycles using the Collabfold_advanced server.

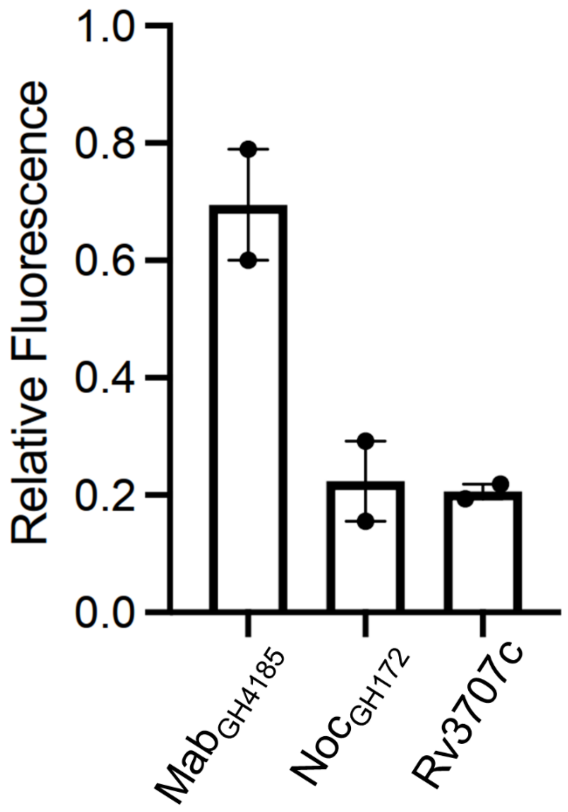


**Figure S16: Release of fluorescently labelled D-arabinan by selected acid-fast enzymes.** Fluorescent mycolyl-arabinogalactan-peptidoglycan complex (mAGP) obtained from 5-AcFPA treated *C. glutamicum* was incubated with 1 μM of each enzyme with rotation overnight. After 3 washes, remaining fluorescent signal was measured and normalized the to an enzyme free control (n =2 technical replicates).

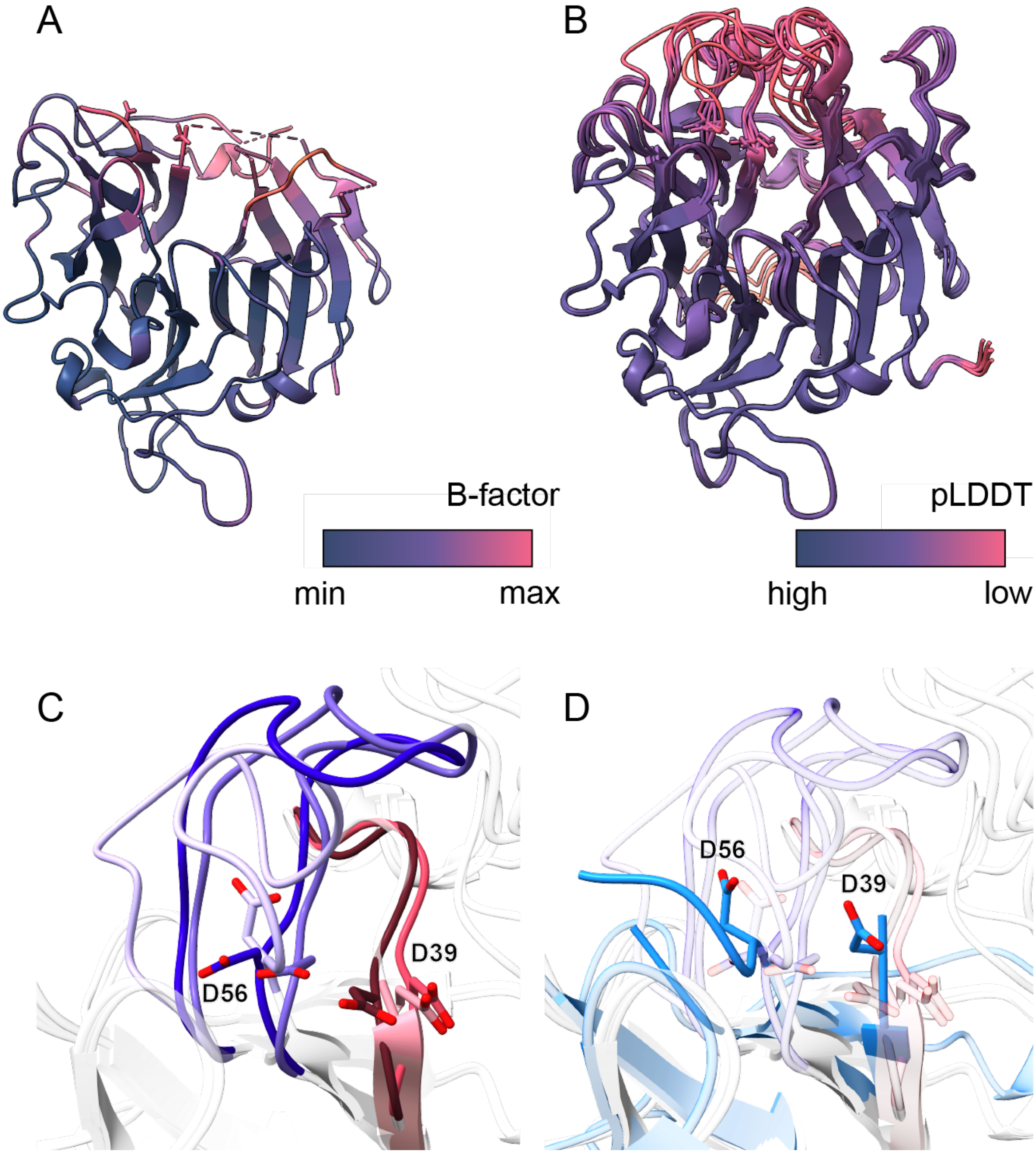


**Figure S17: Multiple catalytic residue conformations are predicted for Rv3707c.** The experimental structure of Rv3707c exhibits significant mobility (**A**) in the same regions where Alphafold has the lowest pLDDT (**B**). The Alphafold-predicted flexibility includes the predicted catalytic residues, the extreme conformations of which are shown in panel **C**. In panel **D** the experimental structure has been overlaid, showing that the experimentally determined catalytic residues are consistent with the Alphafold predictions. In this position the second catalytic residue, D56, is pointed away from the active site suggesting that the protein is in an inactive form.

### Supplementary Tables

**Table S1. Whole cell proteomics of *D. gadei* grown on D-arabinan, ordered by iBAQ**. Proteins from PUL42 are coloured blue.

**Table S2: Data collection and refinement statistics for the crystal structures of Dg$_{GH172c}$ and RV3707c**

| Data statistics* | | |
|---|---|---|
| | **Dg$_{GH172c}$** | **Rv3707c** |
| Beamline | I24 | I04 |
| Date | 12/08/20 | 29/08/21 |
| Wavelength (Å) | 0.9786 | 0.9795 |
| Resolution (Å) | 68.11 - 1.40 (1.42 - 1.40) | 43.50 - 2.41 (2.50 - 2.41) |
| Space group | H32 | H32 |
| Unit-cell parameters | | |
| a (Å) | 89.15 | 145.67 |
| b (Å) | 89.15 | 145.67 |
| c (Å) | 289.32 | 92.68 |
| α = β = γ (°) | 90, 90,120 | 90, 90, 120 |
| Unit-cell volume (Å$^3$) | 1991369 | 1703166 |
| Solvent content (%) | 50.5 % | 47.2% |
| No. of measured reflections | 895646 (28001) | 153262 |
| No. of independent reflections | 87401 (4237) | 14695 (1458) |
| Completeness (%) | 99.9 (99.0) | 99.9 (99.73) |
| Redundancy | 10.2 (6.6) | 10.4 (10.6) |
| CC$_{1/2}$ (%) | 0.99 (0.412) | 1.0 (0.382) |
| I/σ(I) | 12.7 (0.6) | 4.1 (0.62) |
| Refinement statistics* | | |
| Rwork (%) | 14.4 | 21.76 |
| Rfree[#] (%) | 18.3 | 25.70 |
| No. of non-H atoms | | |
| No. of protein, atoms | 3043 | 2265 |
| No. of solvent atoms | 208 | 102 |
| No. of ion atoms | 4 | 12 |
| R.m.s. deviation from ideal values | | |
| Bond lengths (°Å) | 0.019 | 0.005 |
| Bond angles (°) | 2.18 | 1.01 |
| Average B factor (Å$^2$) | | |
| Protein | 26.5 | 57.52 |
| Solvent | 35.0 | 45.93 |
| Ions | 32.4 | 40.64 |
| Ramachandran plot[+], residues in | | |
| Most favoured regions (%) | 96.8 | 95.9 |
| PDB code | 8AH3 | 8AN0 |

*(Values in parenthesis are for the highest resolution shell).
[#]5% of the randomly selected reflections excluded from refinement.
[+]Calculated using MOLPROBITY.

**Table S3: Recipe for 50ml of 2x defined minimal media**

| Component | Mass (mg) or volume (ml) |
|---|---|
| **10x Bacteroides salts (4L)**<br>(544g $KH_2PO_4$, 35g NaCl ,45g $(NH4)_2SO_4$ – pH 7.2) | 10 ml |
| **Balch's vitamins (1L)**<br>(p-Aminobenzoic acid, 5mg<br>Folic acid, 2mg<br>Biotin, 2mg<br>Nicotinic acid, 5mg<br>Calcium pantothenate, 5mg<br>Riboflavin, 5mg<br>Thiamine HCl, 5mg<br>Pyridoxine HCl, 10mg<br>Cyanocobalamin, 0.1mg<br>Thioctic acid, 5mg<br>Distilled Water, 1L) | 1 ml |
| **Trace mineral solution (1L)**<br>EDTA, 0.5g<br>$MgSO_4*7H_2O$, 3g<br>$MnSO_4*H_2O$, 0.5g<br>NaCl, 1g<br>$FeSO_4*7H_2O$, 0.1g<br>$CaCl_2$, 0.1g<br>$ZnSO_4*7H_2O$, 0.1g<br>$CuSO_4*5H_2O$, 0.01g<br>$H_3BO_3$, 0.01g<br>$Na_2MoO_4*2H_2O$, 0.01g<br>$NiCl_2*6H_2O$, 0.02g | 1 ml |
| **Purine/Pyrimidine solution (1L)**<br>200 mg each: Adenine (Sigma, A2786)<br>Guanine (Sigma, G11950)<br>Thymine (Sigma, T0895)<br>Cytosine (Sigma, C3506)<br>Uracil (Sigma, U1128) | 1 ml |
| **Amino acid solution (250 ml)**<br>62.5 mg of all 20 standard amino acids | 1 ml |
| Vitamin K3 (1 mg/ml) | 0.1 ml |
| $FeSO_4$ (0.4 mg/ml) | 0.1 ml |
| $CaCl_2$ (0.8% w/v) | 0.1 ml |
| $MgCl_2$ (0.1 M) | 0.1 ml |
| Vitamin B12 (0.01 mg/ml) | 0.05 ml |
| L-Cysteine | 100 mg |
| Distilled water | Up to 50 ml total |

*Solution pH was adjusted to 7.2 and then syringe-filtered through a 0.22 µM membrane filter. It can then be mixed with any 2x carbon source stock (usually 10 mg/ml) in a 1:1 ratio for a 5 mg/ml final carbon source minimal media.*
